# A pocket-centric framework for selective targeting of amyloid fibril polymorphs

**DOI:** 10.64898/2026.02.25.707901

**Authors:** Gaëtan Ossard, Constantin Bogdan Ciambur, Zachariah P. Schuurs, Ronald Melki, Olivier Sperandio, Eugénie Romero

## Abstract

The rapid expansion of high-resolution cryo-EM structures of amyloid fibrils has transformed our understanding of fibril polymorphism, yet it has not been matched by comparable progress in the rational development of protein-selective or polymorph-specific amyloid ligands. One possible explanation is that ligand selectivity is governed not only by global fibril folds, but also by local surface pockets accessible to small molecules. Here, we present a systematic analysis of 400 cryo-EM structures of amyloid-β, tau, and α-synuclein fibrils. Using a unified pocket similarity index and minimum spanning tree representations, we construct global and protein-specific graph representations of the amyloid binding pocket space, and examine how surface cavities are distributed across proteins, polymorphs, and structural contexts. We find that many detectable pockets are shared across multiple fibrillar folds and, in several cases, across distinct amyloid-forming proteins, suggesting that pocket-level convergence may contribute to the limited selectivity of amyloid-directed ligands. Conversely, only a restricted subset of pockets occupies isolated regions of pocket similarity space, defining rare structural opportunities for protein-selective or polymorph-restricted targeting. Analysis of structures of extracted fibrils further shows that disease-derived fibril pockets do not form a completely isolated pocketome subset, but can resemble pockets observed in selected *in vitro* polymorphs. Together, these results reframe amyloid ligand development as a problem of pocket-level discriminability within a constrained fibril landscape, and provide a structural framework for identifying promising binding sites while avoiding intrinsically non-discriminatory pockets.

**Significance Statement:** Despite major advances in cryo-EM structure determination of amyloid fibrils, the development of selective ligands for amyloid assemblies remains challenging. By systematically comparing surface binding pockets across 400 amyloid-β, tau, and α-synuclein fibrillar structures, we show that many ligand-accessible cavities exhibit similar geometric and physicochemical properties across fibrillar polymorphs made of distinct proteins. This pocket-level convergence provides a structural basis for understanding why many amyloid ligands exhibit broad binding profiles, while also identifying rare pockets that are sufficiently isolated to support more selective targeting strategies. Our work establishes a pocket-centric framework for interpreting amyloid ligand selectivity and for prioritizing fibril binding sites in imaging and therapeutic ligand development.

## Introduction

Alzheimer’s and Parkinson’s diseases are the most prevalent neurodegenerative disorders and represent a growing global health burden. As of 2023, more than 40 million individuals were diagnosed with Alzheimer’s disease (AD), the vast majority of whom were over 60 years of age (1). Clinically, AD is characterized by progressive cognitive decline, while Parkinson’s disease (PD), the most common neurodegenerative movement disorder, presents with motor symptoms such as rigidity, tremor, and bradykinesia, alongside a wide spectrum of non-motor manifestations (2).

Although genetic risk factors have been identified for both disorders (3–5), approximately 90–95% of cases are sporadic, possibly involving environmental stressors (6). At the neuropathological level, AD and PD share key features, including neuroinflammation and the accumulation of proteinaceous aggregates. AD is characterized by extracellular amyloid plaques composed of amyloid-β peptides and intracellular neurofibrillary tangles enriched in tau protein (7). This places AD within the broader class of tauopathies, which also includes chronic traumatic encephalopathy (CTE), frontotemporal dementia (FTD), corticobasal degeneration (CBD), and progressive supranuclear palsy (PSP) (8). PD belongs to the group of synucleinopathies, which includes disorders such as dementia with Lewy bodies (DLB), multiple system atrophy (MSA), and juvenile-onset synucleinopathy (JOS), all linked to the aggregation of α-synuclein into amyloid fibrils that form together with membranous components Lewy bodies and neurites (9, 10).

Importantly, the accumulation of amyloid fibrils in the brain precedes the onset of clinical symptoms by many years, and in some cases by decades, making these aggregates prime targets for early diagnosis and disease monitoring (11, 12). The development of cryo-electron microscopy (cryo-EM) has enabled the determination of amyloid fibril structures at near-atomic resolution, revealing an unexpected degree of structural polymorphism. Both *in vitro* assembled recombinant proteins and fibrils extracted from patient brain tissue have shown that a given protein can adopt distinct fibrillar folds depending on environmental conditions or disease context (13–52). Such different polymorphs were also shown to trigger distinct diseases when delivered to animal models (53–57). In particular, fibrils extracted from patients affected by different tauopathies or synucleinopathies exhibit disease-specific folds, suggesting a structure–pathology relationship (58–63).

Despite this wealth of structural information, the rational development of ligands capable of selectively targeting specific amyloid proteins, or specific fibrillar polymorphs of the same protein, remains highly challenging. Positron emission tomography (PET) has enabled the in vivo detection of amyloid deposits, and several tracers have been approved or advanced in clinical development for amyloid-β and tau imaging (64–66). More recently, α-synuclein tracers such as [¹⁸F]ACI-12589 have shown promise for imaging multiple system atrophy (67). Nevertheless, many amyloid-directed ligands display broad binding profiles across fibril types, limited discrimination between polymorphs, or off-target interactions that restrict their diagnostic and therapeutic utility (68, 69).

This difficulty should not be interpreted as a simple consequence of insufficient structural information. Amyloid ligand design faces several intrinsic challenges: (i) high-resolution fibril structures have only recently become available at scale; (ii) they often lack structural information about flexible regions spanning significant fractions of their constitutive proteins; (iii) the fraction of disease-associated fibrils extracted from patients’ brains whose structures have been solved are often difficult to generate de novo (28, 44); and (iv) the quantitative binding assays for filamentous assemblies remain more complex than assays on soluble globular proteins. Moreover, the few cryo-EM structures of amyloid fibrils in complex with small molecules suggest that ligand recognition can differ substantially from classical binding to a pre-formed protein cavity. In several cases, ligands bind through repeated or stacked arrangements along the fibril axis, combining fibril–ligand and ligand–ligand interactions (33, 48, 49, 70).

Within this broader context, we focus here on one structural determinant that has remained comparatively underexplored: the local surface pockets exposed by amyloid fibrils. While fibril polymorphs are usually classified according to their global fold, small-molecule recognition is governed by the geometry and physicochemical properties of local binding environments. Thus, structurally distinct fibril folds may expose similar ligand-accessible pockets, whereas related folds may present different local binding sites. This raises a central question: does amyloid fibril polymorphism generate sufficiently distinct pocket landscapes to support protein-selective or polymorph-specific ligand design?

To address this question, we adopt a pocket-centric perspective and perform a systematic pocketome analysis of amyloid fibrils. Pocketomes were originally developed to catalogue and compare ligand-binding cavities in proteins and protein–protein interaction interfaces (71–74), providing a quantitative framework to compare binding environments independently of global fold similarity. Here, we extend this concept to amyloid assemblies by analyzing surface pockets across all cryo-EM structures of amyloid-β, tau, and α-synuclein fibrils available in the Protein Data Bank as of February 2026. Importantly, our aim is not to assign equal biological relevance to all *in vitro*, seeded, mutant, extracted and ligand-bound structures, but to map the structural envelope of pocket diversity across experimentally observed fibrillar states. Disease-derived structures are then analyzed explicitly to determine whether their pockets occupy a distinct region of the amyloid pocketome or overlap with pockets observed in other fibrillar contexts.

By detecting surface-accessible pockets, comparing them through a unified pocket similarity index, and visualizing their relationships using minimum spanning tree representations, we construct global and protein-specific amyloid pocketomes. This framework enables us to identify broadly recurrent pockets that are likely to support non-discriminatory binding, as well as rare isolated pockets that may offer more realistic opportunities for selective targeting. In doing so, we reframe amyloid ligand development not as a direct consequence of fibril fold classification, but as a problem of pocket-level discriminability within a constrained landscape of amyloid surface cavities.

## Results

### Detection of surface-accessible pockets across amyloid fibril polymorphs

To systematically compare ligand-relevant binding environments across amyloid fibrils, we first established a unified workflow for structure selection, pocket detection and pocket filtering. We retrieved 400 cryo-EM structures of α-synuclein, tau, and amyloid-β fibrils available in the Protein Data Bank as of February 2026, including *in vitro* assembled fibrils, filaments purified from patients’ brain homogenates, and seed amplification assay products (Fig. 2, Fig. S1-S3 and Supplementary Files 1-3).

**Figure 1.**
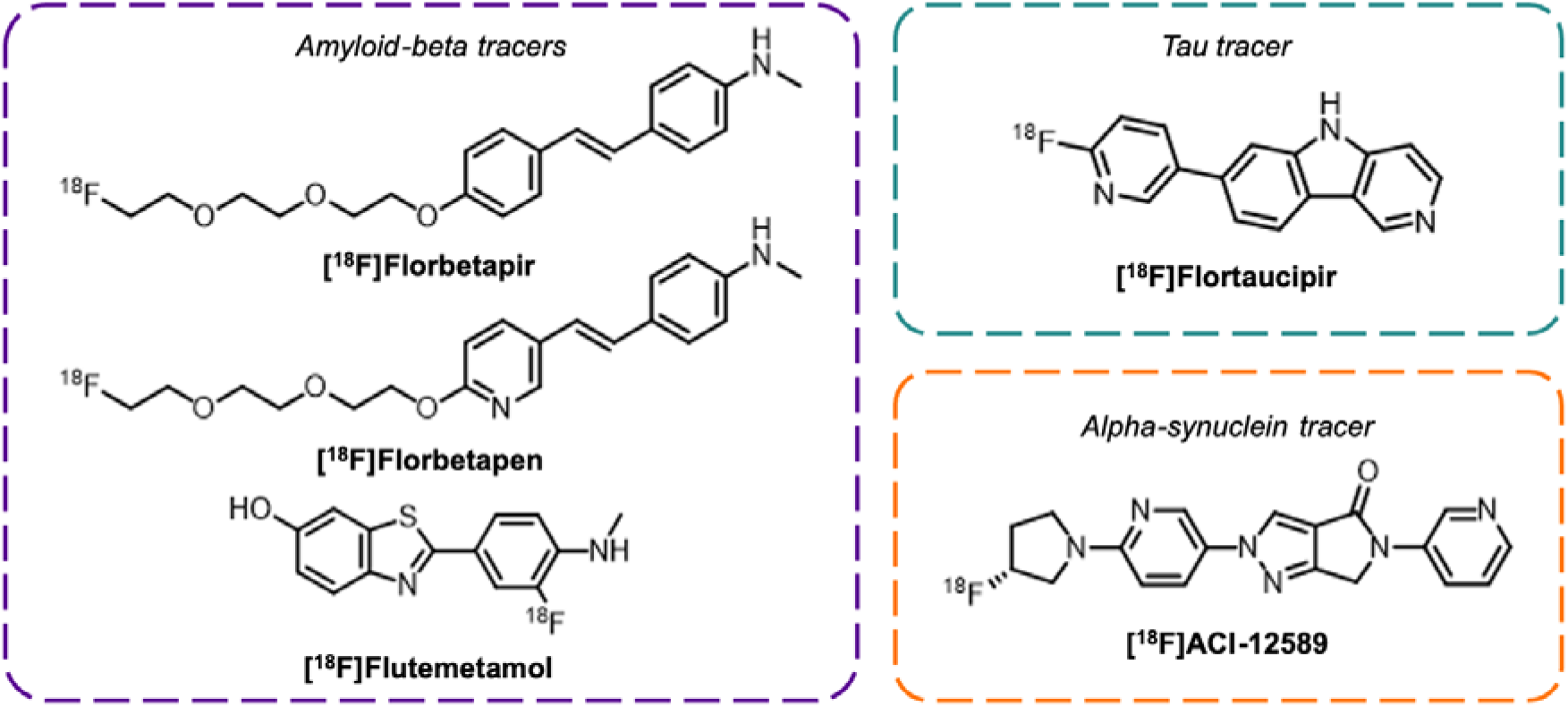
Clinically-validated amyloid PET tracers for imaging amyloid-β (purple), tau (blue) and α-synuclein (orange) fibrils

**Figure 2.**
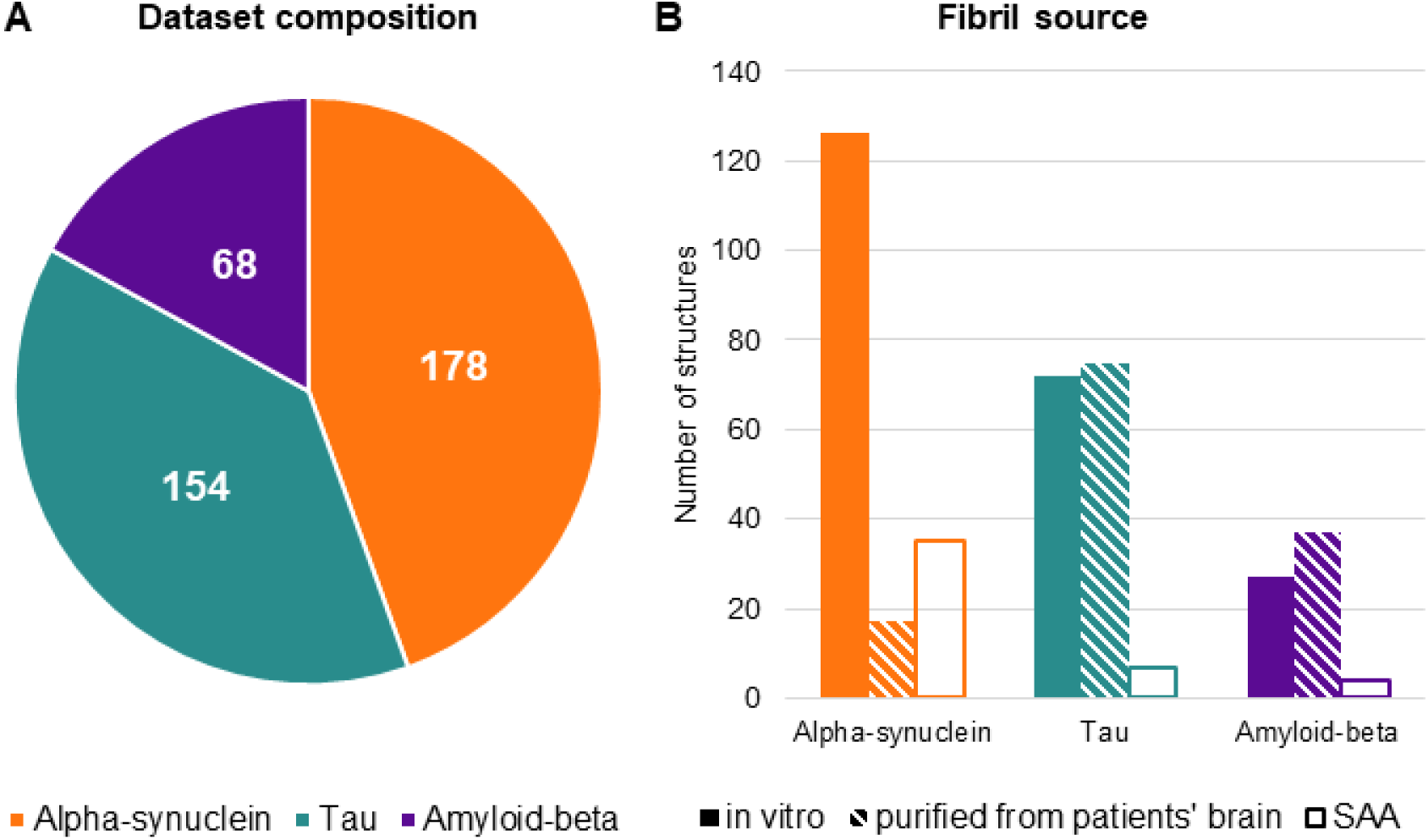
Detailed analysis of the dataset. A) Number of structures in the dataset. B) Method used to obtain the fibril. SAA = Seed Amplification Assay.

Because many deposited fibril structures correspond to closely related or identical folds, analyzing each entry independently would introduce substantial redundancy into the pocketome. We therefore classified structures within each amyloid protein family using the Calypso workflow developed by Connor et al., based on fold-level RMSD relationships (52, 75). Distance thresholds were selected to define a tractable set of representative polymorph groups for each protein, yielding approximately 15 groups per amyloid family (Fig. 3; Fig. S4–S5). These groupings were broadly consistent with previously reported classifications of amyloid fibril polymorphs (52, 76–78). For each group, we selected the best-resolved representative structure for downstream pocket analysis (Fig. S6–S8). Two α-synuclein entries, 7YK8 and 9JI8, were excluded because their short, flat, and partially unresolved peptide segments did not provide suitable fibrillar models for pocket detection. For Pick’s disease tau filaments, 6GX5 was retained as the representative structure because the better-resolved entry 8P34 contains only a single chain in the deposited PDB model, precluding reliable pocket detection on a fibrillar assembly.

**Figure 3.**
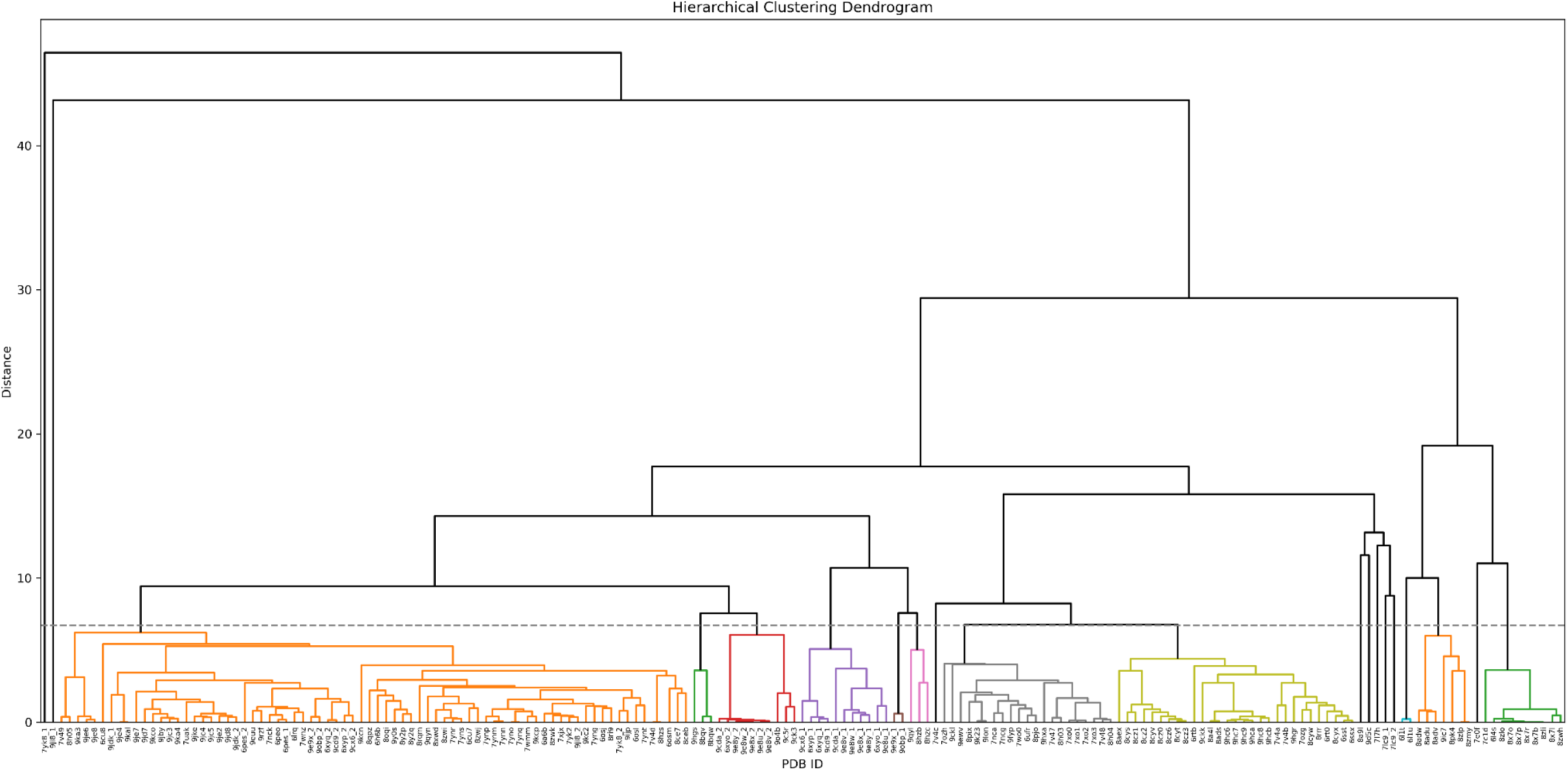
Hierarchical classification of α-synuclein fibrils. The procedure was performed according to Connor et al. (52, 75). Fibrils were classified into 18 oups, cutting at a distance of 5.3 Å. Fibrils from same groups are colored accordingly.

Surface-accessible cavities were then detected on each representative fibril using VolSite (79) (Fig. 4A–C). Because amyloid fibrils contain grooves, inter-protofilament interfaces, buried channels, and fibril-end cavities that are not all equally relevant to small-molecule recognition, we applied stringent filtering criteria to retain only pockets that are exposed at the fibril surface and repeated along the fibril axis. Cavities confined to fibril tips, buried within the fibril core, located at protofilament interfaces, too small, or geometrically constrained were excluded (Fig. 4C; Fig. S9). This procedure was designed to focus the analysis on recurring surface environments that could plausibly contribute to ligand binding, rather than on inaccessible or artefactual cavities.

**Figure 4.**
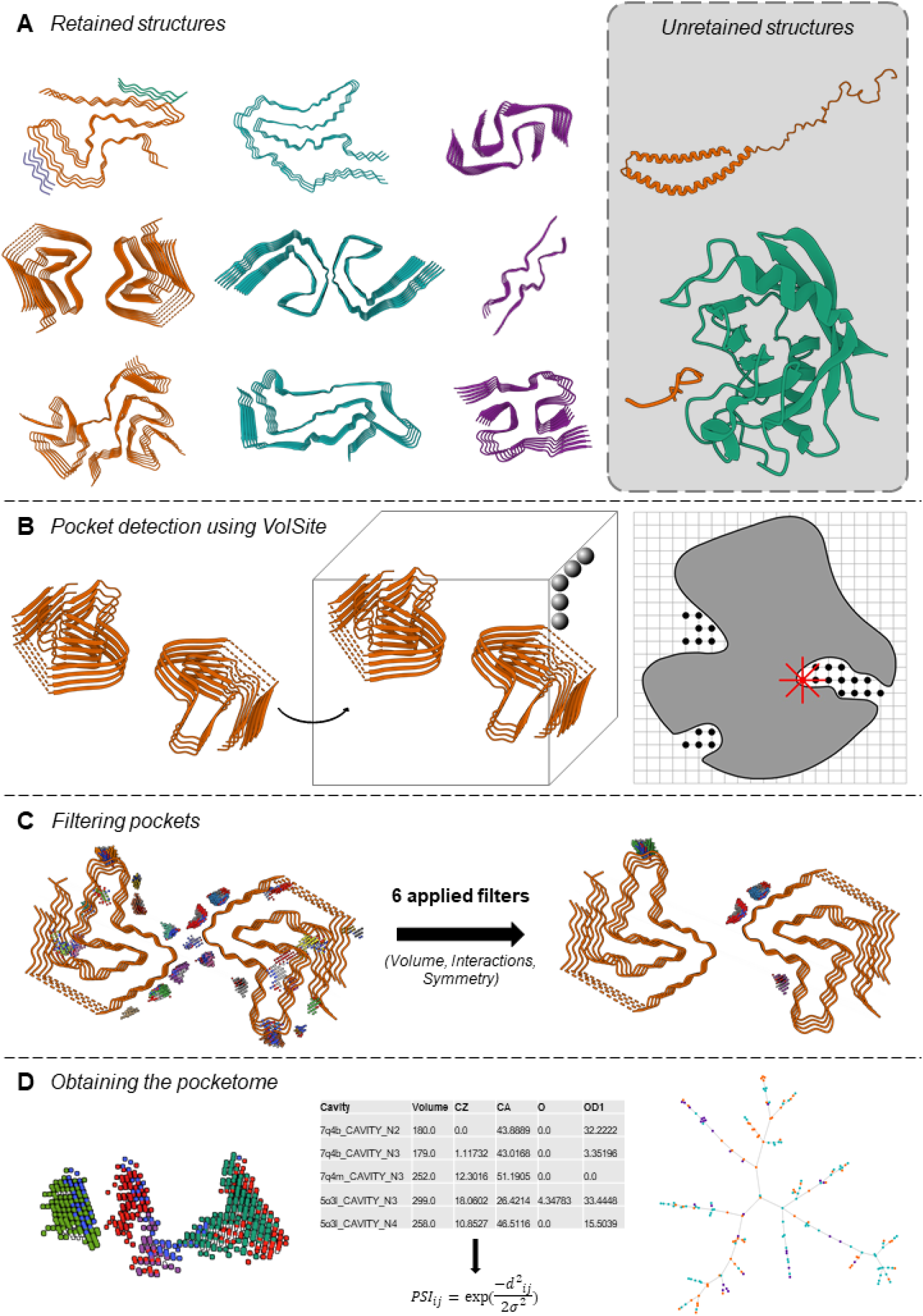
Overview of the presented study. A) Examples of fibril structures of α-synuclein (orange), tau (blue) and amyloid-β (purple) retrieved from the PDB. Examples of non-considered structures for the accession code SNCA are shown in the grey area, with α-synuclein shown in orange. PDB IDs from the shown examples (top to bottom, left to right): 8A9L, 6SSX, 6XYQ, 6TJO, 5O3L, 7P65, 9FH2, 8FF3, 7Q4B (retained structures), 1XQ8 and 6I42 (unretained structures). B) Schematic of the pocket detection step using α-synuclein representative entry (PDB ID: 9CKK) from polymorph group 8 as an example (left image). The protein is protonated and then placed in a three-dimensional matrix (represented as a cube, middle image), that is composed of regularly spaced spheres. VolSite algorithm iteratively goes through each sphere, projecting rays (shown in red) and retrieving interaction to determine the final state of the sphere (right image, modified from Weisel et al. (84)). C) Results of pocket detection before and after application of all six considered filters on α-synuclein polymorph group 8 fibril. D) Generation of the pocketome and visualization using a minimum spanning tree approach. The collection of pockets corresponds to a table representing each pocket using geometric descriptors (5 shown as example in the table). The non-zero descriptors are used to calculate pairwise pocket similarity indexes that are then used to create the graph and visualize the pocketome.

As an internal validation of this detection strategy, we also applied the same VolSite procedure to available cryo-EM structures of ligand-bound amyloid fibrils. The detected cavities overlapped well with the experimentally reported ligand-binding regions described in the corresponding structural studies (Fig. S10), supporting the relevance of the filtered pockets as ligand-accessible surface environments.

The resulting set of pockets constitutes the fundamental unit of the amyloid pocketome. Each retained pocket was encoded using a common descriptor space capturing geometric and physicochemical properties, enabling quantitative comparison of binding environments across proteins, polymorphs, and structural contexts (Fig. 4D).

### The global amyloid pocketome reveals extensive cross-protein pocket similarity

We next asked whether amyloid fibrils formed by different proteins expose distinct or overlapping ligand-accessible pocket landscapes. To address this question, all retained surface pockets from amyloid-β, tau, and α-synuclein fibrils were compared within a unified descriptor space. Pairwise pocket similarity was quantified using the Pocket Similarity Index (PSI) (Equation 1), and the resulting relationships were visualized as a minimum spanning tree in which each node represents a pocket, and neighboring nodes correspond to locally similar binding environments (Fig. 5).

**Figure 5.**
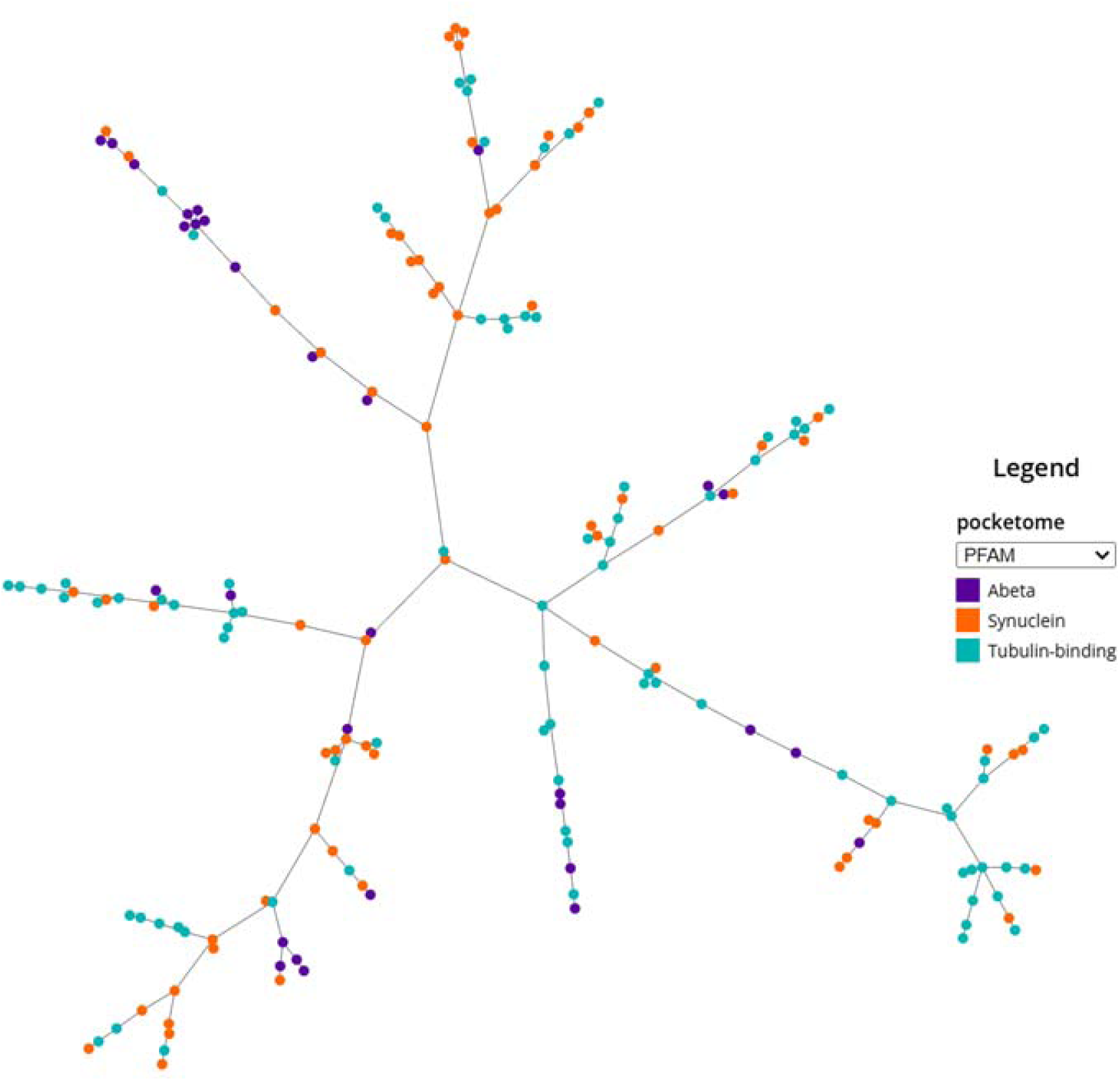
The Amyloid pocketome, represented as a minimum spanning tree. Each pocket is represented using a dot, and dots are connected using the PSI values, so that the total number of branches is minimal. The legend shown here represents fibril type, where dots are colored according to the three different proteins, amyloid-β (purple), α-synuclein (orange) or tau (blue).

If protein identity were the dominant determinant of pocket architecture, pockets from amyloid-β, tau, and α-synuclein would be expected to occupy distinct regions of the tree. Instead, the global amyloid pocketome forms a largely continuous landscape in which pockets from the three amyloid-forming proteins are extensively intermingled. Although local enrichments of protein-associated pockets can be observed, these regions do not form globally isolated branches. Rather, they remain embedded within a shared pocket similarity space populated by pockets from multiple amyloid proteins.

This organization indicates that amyloid fibrils with distinct sequences and global folds can nevertheless expose locally similar surface cavities. Thus, protein identity and fibril fold are not sufficient to define ligand-relevant binding environments. At the pocket level, many amyloid fibrils converge toward similar combinations of size, shape, accessibility, and physicochemical properties. This convergence is particularly important because small-molecule recognition is governed by the local environment encountered by the ligand, rather than by the overall fibril architecture.

The extensive intermixing observed in the global pocketome provides a structural rationale for the broad binding profiles frequently reported for amyloid-directed ligands. Pockets located within densely populated regions of the tree are not unique to a single amyloid protein, and ligands targeting such sites would be expected to display limited protein selectivity. Conversely, only pockets occupying more isolated regions of the pocketome would be expected to offer realistic opportunities for selective ligand recognition. These results therefore establish pocket-level convergence as a major structural constraint on amyloid ligand selectivity.

### Pocket similarity is decoupled from fibril fold and polymorph classification

We next examined whether pocket similarity follows the polymorph classification derived from fibril fold comparisons. When the global amyloid pocketome was colored by polymorph group identity, pockets from the same fold-defined group did not systematically co-localize within the tree (Fig. S11). Instead, pockets from distinct polymorphs frequently occupied neighboring positions, whereas different pockets from the same polymorph were often distributed across distant branches. Thus, a fibril polymorph does not correspond to a single, homogeneous pocket environment, but rather exposes multiple surface cavities with distinct similarity relationships.

To determine whether this apparent decoupling persists within individual amyloid protein families, we generated protein-specific pocketomes for α-synuclein, tau, and amyloid-β (Fig. 6; Fig. S12–S13). This analysis directly addresses whether polymorph-restricted pocket environments emerge once cross-protein diversity is removed. Across all three proteins, the pocketomes remained heterogeneous and only weakly structured by fold-defined polymorph identity. In the α-synuclein pocketome, pockets from the different polymorph groups were broadly distributed throughout the tree, with no dominant branches corresponding to individual polymorphs (Fig. 6). Similar patterns were observed for tau and amyloid-β, including disease-associated folds, whose pockets remained interspersed with pockets from other structural contexts rather than forming isolated regions (Fig. S12–S13).

**Figure 6.**
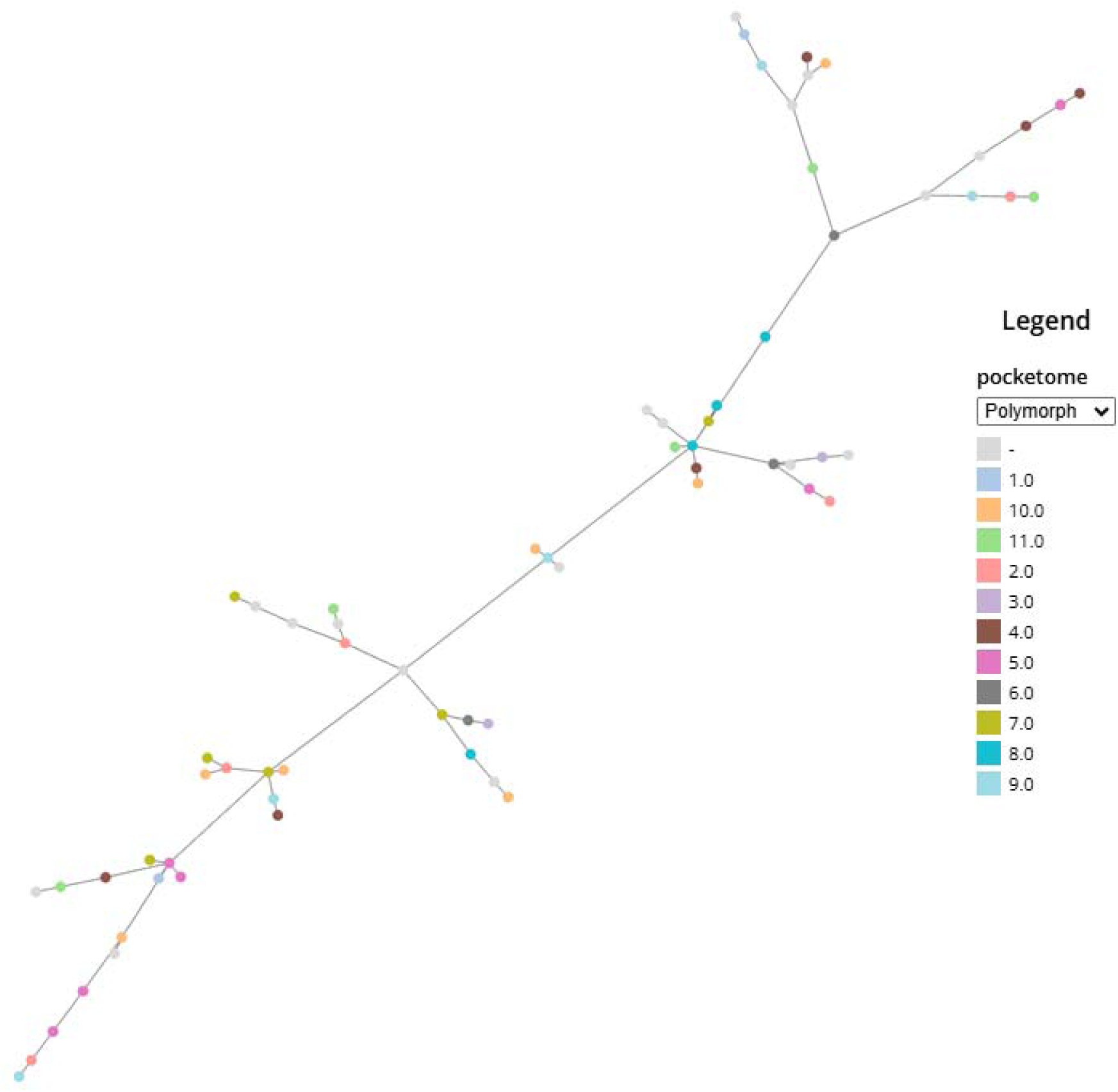
The α-synuclein pocketome, represented as a minimum spanning tree. Each pocket is represented using a dot, and dots are connected using the PSI values, so that the total number of branches is minimal. The legend shown here represents pockets’ volume.

These observations demonstrate that pocket similarity is not a direct consequence of fibril fold similarity. Structurally distinct polymorphs can expose locally similar surface pockets, while pockets from the same polymorph can differ substantially from one another. This decoupling has important implications for amyloid ligand design. A ligand selected against one pocket of a given fibril polymorph may recognize similar pockets present in other polymorphs, whereas other pockets on the same fibril may not be recognized at all. As a result, polymorph-specific targeting cannot be inferred from fold classification alone.

### Recurrent sequence motifs underlie non-discriminatory pocket classes

To better understand why many pockets occupy shared regions of pocketome space, we annotated each pocket according to the amino-acid residues contributing to its boundaries. This residue-level analysis showed that recurrent pocket classes are often defined by short local motifs rather than by the global architecture of the fibril. In particular, highly connected regions of the pocketome frequently correspond to shallow cavities formed by neighboring charged, polar, or aromatic residues, including lysines, arginines, glutamines, and tyrosines.

In α-synuclein, several recurrent pocket classes are formed between closely spaced basic residues, including K21–K23, K43–K45, and K58–K60. These sites define small and elongated surface cavities, typically with volumes in the range of 100–150 Å³. Their chemical composition may provide favorable electrostatic or hydrogen-bonding opportunities, but their limited size and shallow geometry impose strong constraints on ligand accommodation. Such pockets are therefore unlikely to support highly selective recognition by a single small molecule unless additional interactions, such as repeated binding along the fibril axis or ligand–ligand stacking, contribute to binding.

This residue-level view explains why pocket similarity can be partly decoupled from fibril fold similarity. Distinct fibril architectures can repeatedly expose comparable local arrangements of side chains, generating pockets that appear similar to a ligand even when the underlying backbone folds differ. Conversely, different surface regions of the same fibril can generate chemically and geometrically distinct pockets. Thus, pocket recurrence arises from local residue environments rather than from polymorph identity alone.

These recurrent local motifs represent structurally plausible but poorly discriminating binding environments. Their abundance across the pocketome suggests that many amyloid-directed ligands may bind broadly, not because they recognize a conserved global fibril fold, but because they engage common local surface motifs exposed by multiple amyloid assemblies. Identifying such motifs is therefore useful not only for explaining off-target binding, but also for deprioritizing pockets that are intrinsically unlikely to support protein-selective or polymorph-specific ligand design.

### Pocketome clustering identifies rare isolated pockets compatible with selective targeting

Having established that most amyloid pockets occupy a continuous and partially overlapping similarity landscape, we next sought to identify the unique pockets that might support selective recognition. We reasoned that a pocket compatible with protein-selective or polymorph-restricted ligand design should satisfy two conditions: it should be defined by a unique binding environment, and it should be globally isolated from unrelated pockets, so that ligands targeting it are less likely to cross-react with other amyloid assemblies/polymorphs.

To identify such environments, we applied density-based clustering to the PSI-derived distance matrices obtained from the global and protein-specific pocketomes. This analysis was not intended to assign biological function to individual pockets, but to distinguish compact and structurally discriminable pocket classes from the continuous background of weakly separated surface cavities (Fig. S14). Across the datasets, most pockets were not assigned to any cluster, consistent with the largely continuous organization of amyloid pocket similarity space. In contrast, a limited number of compact clusters were detected, corresponding to recurrent pocket environments that remain relatively isolated from the rest of the pocketome.

These isolated clusters fall into two main categories. Within the first, protein-conserved pockets, largely restricted to the amyloid-forming protein, are shared by several polymorphs of the considered protein. A representative example is amyloid cluster 6, composed exclusively of tau pockets (Fig. S15A). This cluster includes pockets from tau filaments extracted from Pick’s disease (6GX5), tau P301L and P301T mutant fibrils (9GG0 and 9GG1), CBD type II tau filaments (6TJX), CBD-seeded fibrils (8ORG), and *in vitro* fibrils formed from a partial tau construct in the presence of KCl (7R5H). Despite differences in global fibril fold, disease context, and assembly conditions, these pockets retain a similar local geometry, reflecting conserved side-chain arrangements. Such pockets may represent plausible targets for protein-selective but polymorph-agnostic ligands.

The second category, termed “cross-amyloid pockets”, corresponds to structurally similar cavities found in different amyloid-forming proteins (Fig. S15B). Amyloid cluster 4, which includes small pockets of approximately 150 Å³ or less from extracted α-synuclein JOS fibrils (8BQV), limbic-predominant neuronal inclusion body 4R tauopathy type 1a tau filaments (7P6A), and AD-PHF-like *in vitro* tau fibrils (8Q8R) perfectly represents this category of pockets. These pockets are shallow, geometrically constrained, and chemically simple, often formed around adjacent lysine, arginine, glutamine, or glycine residues. Their recurrence across unrelated amyloid proteins identifies them as intrinsically non-discriminatory environments. Although such pockets may support ligand binding, they are poor candidates for selective ligand design because structurally similar binding sites exist elsewhere in the amyloid pocketome.

Notably, we did not identify compact clusters composed exclusively of multiple pockets from a given polymorph. This suggests that polymorph-specific binding environments are rare in the current structural dataset. However, some individual pockets occupy terminal positions in the pocketome tree and display very low similarity to the rest of the dataset, with PSI values below 0.005. These pockets represent structural outliers rather than recurrent clusters. They are often large or more complex cavities, defined by four or more contributing residues, and may therefore provide the most plausible starting points for polymorph-restricted ligand design (Fig. S15C). At the same time, their uniqueness also implies a more challenging ligand-design problem, because such pockets may be less repetitive, less shallow, and less chemically simple than the recurrent non-discriminatory pockets.

Together, these analyses show that selective amyloid targeting is governed not by the mere presence of detectable pockets, but by their position within the global pocketome. Pockets embedded in densely populated regions are intrinsically prone to cross-reactivity, whereas isolated clusters or extreme outliers define the limited subset of binding environments for which selective recognition may be structurally plausible.

### Overlap between pockets within α-synuclein fibrils extracted from patients’ brains and selected *in vitro* assembled fibrils

We next focused on α-synuclein fibrils, for which the relationship between *in vitro* assembly models and disease-derived structures remains a central issue for ligand discovery. Unlike several tau fibril folds, which have been reproduced *in vitro* (28) or by seeding from disease-derived material, α-synuclein fibrils extracted from the brain of patients developing distinct synucleinopathies remain difficult to recapitulate experimentally (44). Numerous *in vitro* assembly conditions generate distinct α-synuclein polymorphs, some of which induce very efficiently pathological phenotypes in cellular or animal models (53, 54), but their folds differ from those reported for fibrils extracted from the brain of a very limited number of MSA, PD, PDD patients. We therefore asked whether the latter fibrils expose pockets that are structurally distinct from all *in vitro* models, or whether selected *in vitro* polymorphs contain comparable binding environments.

The α-synuclein pocketome showed that disease-derived pockets do not form a completely isolated subset. MSA fibrils are composed of two distinct protofilaments, both of which fall within larger structural groups in the fold-based classification, whereas the PD/PDD/DLB Lewy fold is represented by a single protofilament that is more distinct from the rest of the dataset (Fig. 3). At the pocket level, however, several cavities within extracted fibrils were found to resemble pockets from selected *in vitro* fibrils.

Two pockets from MSA type I protofilament IA illustrate this relationship (Fig. 7A). The first is a small N-terminal pocket enriched in lysine residues, corresponding to pocket 15, which belongs to α-synuclein cluster 1 together with pockets from *in vitro* fibrils formed in the presence of ATP (9QYL) or induced by heparin (7V4C). This MSA pocket showed close similarity to 7V4C pocket 7, with a PSI value of 0.624. A second, larger N-terminal pocket formed between G14 and K32 was also related to an *in vitro* ligand-binding environment, showing similarity to the F0502B-binding cavity observed in WT-polymorph 5A fibrils (8ZMY), with a PSI value of 0.335. These observations suggest that selected *in vitro* α-synuclein polymorphs can reproduce local binding environments that resemble pockets present in disease-derived MSA fibrils, even when their global fibril folds are not identical.

**Figure 7.**
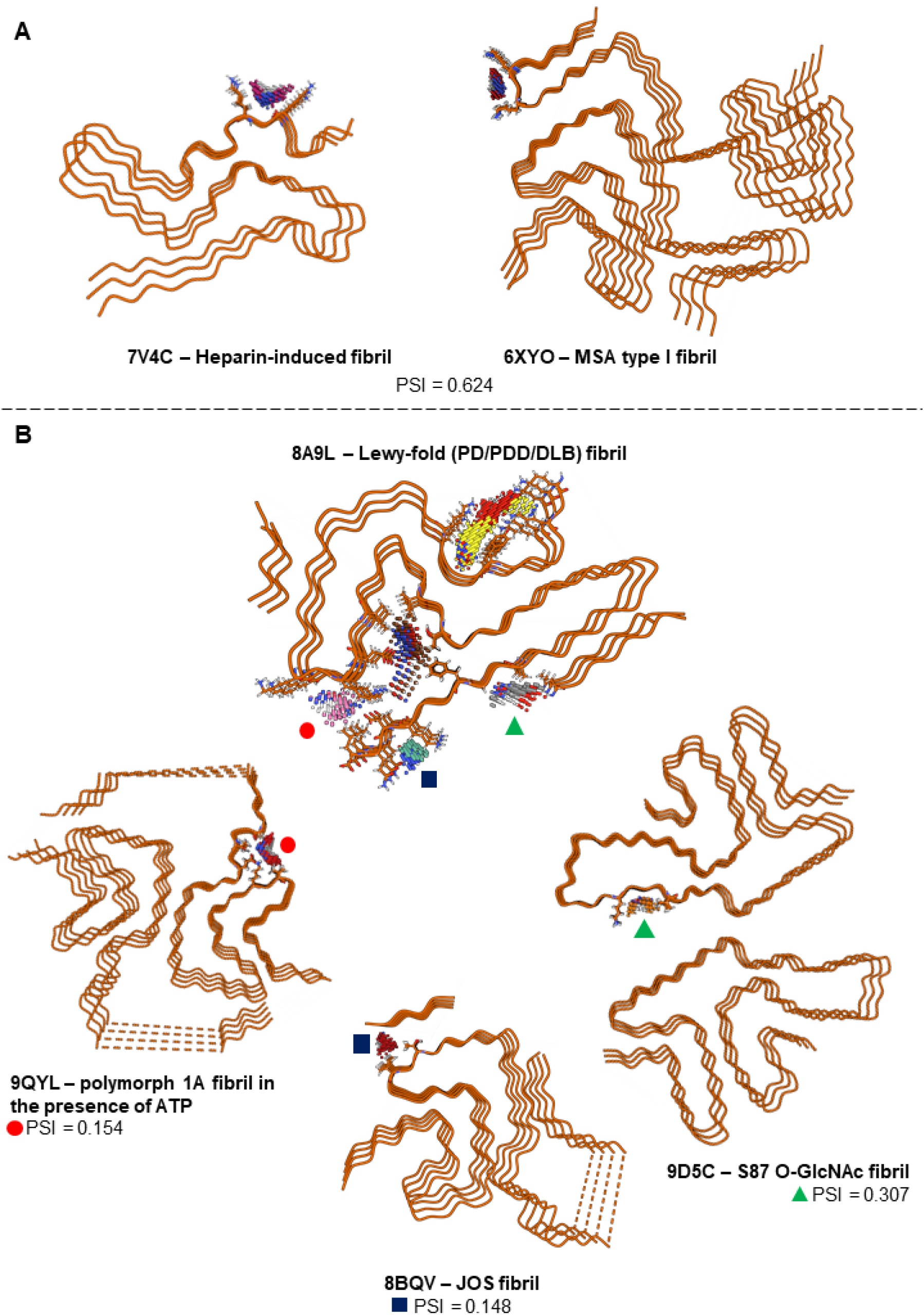
Pockets from patient-derived fibril structures show structural similarities with other polymorphs. A) Pocket number 15 from MSA-type I fibrils (PDB: 6XYO) was found highly similar to pocket number 9 from heparin-induced recombinant fibrils (7V4C), with a pairwise PSI value of 0.624. B) Pockets from extracted fibrils found in patients with Parkinson’s disease, Parkinson’s derived dementia and Dementia with Lewy bodies adopting the Lewy fold (8A9L) show similarities with other pockets. Pocket number 5 (red dot) is similar with pocket number 7 from recombinant fibrils formed in the presence of ATP (9QYL), with a PSI value of 0.154. Pocket number 10 (blue square) is similar with pocket number 7 from Juvenile-onset synucleinopathy fibrils (8BQV), with a 0.148 PSI value. Pocket number 6 (green triangle) is similar with pocket number 9 from Ser87-glycosylated recombinant fibrils (9D5C), with a PSI value of 0.307. The two largest pockets (number 1 and 3) from the Lewy-fold fibrils appear to be unique, with PSI values of 0.005 to their first neighbor. Contributing pocket residues’ side chains are highlighted and pockets represented as dense point clouds.

The situation is more nuanced for fibrils from PD/PDD/DLB patients (Fig. 7B). The two largest cavities detected in the PDD structure displayed highly distinctive geometries, with PSI values of 0.005 or lower relative to the rest of the pocketome. These pockets represent potential structural outliers, but should be interpreted cautiously as ligand-design opportunities. The largest cavity, located between K32 and K45, coincides with an unidentified density reported in the original cryo-EM map, while the second largest pocket is comparatively buried and poorly accessible. In contrast, the remaining smaller pockets from structures from PD/PDD/DLB patients were closer to pockets observed in other α-synuclein polymorphs.

Together, these results show that pockets within patient-derived α-synuclein fibrils are neither completely distinct from, nor broadly shared with, *in vitro* fibrillar polymorphs.. Some disease-derived pockets overlap with binding environments found in selected *in vitro* fibrils, supporting the use of carefully chosen *in vitro* polymorphs as structural models for ligand discovery. At the same time, isolated pockets within fibrils extracted from patients may provide opportunities for more selective targeting, but only if their accessibility, repeatability along the fibril axis, and compatibility with ligand binding can be established experimentally. This pocket-centric view therefore provides a more nuanced framework than global fold comparison alone for selecting α-synuclein fibril models in ligand-design studies.

## Discussion

### Amyloid ligand selectivity is constrained by pocket-level convergence

The rapid expansion of high-resolution amyloid fibril structures has profoundly changed our understanding of fibril polymorphism and disease-associated amyloid strains (62). However, this structural progress has not yet translated into a comparable ability to design ligands that selectively recognize a given amyloid protein, disease-associated fold, or fibril polymorph (69). The present study provides a structural explanation for this disconnect. By comparing ligand-accessible surface pockets across amyloid-β, tau, and α-synuclein fibrils, we show that local binding environments are far less diverse than the fibril folds that display them.

The main conclusion of this work is that amyloid polymorphism does not automatically generate pocket-level discriminability. Fibrils that differ in sequence, global architecture, protofilament arrangement, or disease context can nevertheless expose surface cavities with similar size, shape, accessibility, and physicochemical properties. Conversely, different pockets on the same fibril can occupy distant regions of pocket similarity space. Thus, the structural unit that matters for ligand recognition is not the fibril fold as a whole, but the local pocket accessible to the ligand.

This observation reframes a long-standing problem in amyloid ligand development. Broad binding profiles and off-target recognition are often interpreted primarily as limitations of ligand chemistry. Our results suggest that they can also arise from the pocket landscape itself. If a ligand targets a cavity embedded in a densely populated region of amyloid pocketome space, structurally similar binding environments are likely to exist on other proteins or polymorphs. In such cases, improving ligand affinity may not be sufficient to achieve selectivity, because the underlying pocket is intrinsically non-discriminatory.

### Recurrent local environments explain why specificity is rare

The residue-level analysis helps explain why pocket-level convergence is so widespread. Many recurrent pockets are built from short local arrangements of charged, polar, or aromatic residues that are repeatedly exposed on fibril surfaces. These motifs can generate chemically attractive binding environments, but they are often shallow, small, and geometrically constrained. Such pockets may support amyloid ligand binding, especially when binding is reinforced by repetition along the fibril axis or by ligand–ligand stacking, but they provide limited structural information for discriminating between related amyloid assemblies.

This distinction is important because a pocket can be both suited for binding and poorly selective. In classical structure-based ligand design, the presence of a well-defined cavity is often taken as a favorable starting point. In amyloid fibrils however, the relevant question is not only whether a pocket can accommodate a ligand, but whether that pocket is sufficiently distinct from the rest of the amyloid pocketome. A shallow lysine- or glutamine-rich groove may be chemically accessible, yet if comparable grooves occur repeatedly across unrelated fibrils, ligands directed toward such sites are expected to display broad recognition profiles.

The rare pockets that appear most compatible with selective targeting are therefore not necessarily the most obvious or abundant cavities. Rather, they are pockets that are isolated in pocket similarity space, either as compact protein-restricted clusters or as individual structural outliers. Protein-conserved pockets may be useful for designing ligands that recognize a class of fibrils, such as tau or α-synuclein assemblies, without distinguishing every polymorph. In contrast, polymorph-restricted targeting would require pockets that are not only accessible and repeated along the fibril axis but also sufficiently distinct from all other amyloid pockets. Our analysis suggests that such pockets are uncommon in the current structural dataset. The reason behind this finding is unclear and may be related to fibrils’ helicity that constrains key residues’ folding events. Fibrillar polymorphs lacking helical symmetry, such as ribbons (13), are not represented within the PDB and could well help to identify unique binding opportunities.

### Implications for *in vitro* models, extracted fibrils, and ligand discovery

The pocketome framework also provides a practical way to reassess the value of *in vitro* fibril models for amyloid ligand discovery. *In vitro* assemblies are often considered imperfect models of disease because their global folds can differ from those extracted from patients. This concern is particularly acute for α-synuclein, where disease-derived fibrils remain difficult to reproduce experimentally. Our results suggest that the relevance of an *in vitro* model should not be judged only by global fold similarity, but also by whether it reproduces the local pocket environment that a ligand is intended to target.

This distinction is important for structure-guided ligand design. An *in vitro* fibril that does not match an extracted fold globally may still expose a pocket comparable to a disease-derived binding site and may therefore serve as a useful model for studying ligand recognition. Conversely, an *in vitro* fibril that appears globally related to a disease-associated structure may expose a different set of surface cavities and may be less informative for ligand optimization. A pocket-centric comparison can therefore guide the selection of experimental fibril models by matching the local binding environment rather than relying exclusively on fold classification.

Analysis of fibrillar α-synuclein polymorphs illustrates this point. Some pockets detected in fibrils extracted from MSA patients overlap with pockets found in selected *in vitro* polymorphs, including ATP- or heparin-associated fibrils and ligand-bound WT-polymorph 5A fibrils. These relationships do not imply that the *in vitro* structures reproduce the full disease-associated fibril fold. Rather, they indicate that specific local binding environments can be shared between *in vitro* and extracted polymorphs. Such overlap may justify the use of selected *in vitro* fibrils as tractable structural models for ligand discovery, provided that the targeted pocket is the relevant shared feature.

At the same time, the pocketome highlights pockets that should be deprioritized. Cross-amyloid pockets, especially small and shallow cavities formed by recurrent local motifs, may be attractive from a purely chemical perspective but are intrinsically prone to off-target binding. For ligand development, excluding such non-discriminatory pockets may be as important as identifying promising ones. In this sense, the amyloid pocketome can function as a negative-design tool: it helps identify binding environments where selectivity is unlikely to emerge, even after extensive chemical optimization.

Such observations partially match the conclusions of Sadek et al. (preprint, 80), who developed a computational pipeline entitled FibrilSite to identify interaction sites at the surface of amyloid fibrils, compare their properties and determine the most druggable sites for ligand development prioritization. The method we present here and that by Sadek et al., shows overlap, in particular for fibrillar α-synuclein polymorphs made *in vitro* or extracted from MSA patients. However, fewer but larger sites were identified by the latter method, leading to the conclusion that no similar sites exist between amyloid-forming proteins, in contradiction with the observed lack of selectivity of available and developed ligands.

For diagnostic imaging, the desired selectivity profile will depend on the intended application. Despite limiting the design of ligands that discriminate between closely related pathological strains, the presence of pockets shared across multiple polymorphs is of interest. Such pockets allow the binding of ligands targeting a broader class of assemblies, such as pan-synucleinopathy or pan-tauopathy tracers and/or co-pathologies such as those observed in DLB or PSP and brain region-based disease diagnosis and follow-up. In contrast, tracers intended to distinguish disease subtypes or fibril strains would require pockets that are both accessible and isolated from the broader pocketome. The present analysis suggests that the latter case is structurally much more constrained and will require careful prioritization of rare pocket outliers rather than broadly recurrent cavities. These considerations therefore reinforce the need to evaluate amyloid targets at the level of pocket discriminability rather than at the level of fibril fold identity alone.

### Limitations and outlook

Several limitations should be considered when interpreting this pocketome analysis. First, the detection of a surface pocket does not demonstrate ligand binding, high affinity, or selectivity. The pockets identified here are structural environments that are accessible and repeated along fibril surfaces, but their functional relevance must be established experimentally. Pocket isolation in similarity space should be viewed as a prioritization criterion rather than as a direct prediction of ligandability.

Second, amyloid ligand binding can differ substantially from classical small-molecule recognition by globular proteins. In several cryo-EM structures of ligand-bound amyloid fibrils, ligands bind in repeated arrays along the fibril axis and may be stabilized by both fibril–ligand and ligand–ligand interactions. Such cooperative or stacking effects are not fully captured by pocket descriptors based on local cavity geometry and physicochemical properties. As a result, the pocketome provides information on the local binding environment, but not on the full thermodynamic or supramolecular binding mechanism.

Third, amyloid fibril structures usually resolve only the ordered fibrillar core. Flexible N- or C-terminal regions, post-translational modifications, cofactors, lipids, and other cellular components may contribute to ligand recognition in vivo but are absent or incompletely represented in most structural models. This is particularly relevant for α-synuclein and tau, whose disordered regions may influence accessibility, surface chemistry, interactions with other molecules or even define additional unforeseen binding sites. Future analyses integrating cryo-EM density, molecular dynamics, and experimentally mapped ligand-binding data will be needed to refine the relationship between detected pockets and true binding sites.

Finally, the pocketome is limited by the structural diversity available in the Protein Data Bank at the time of analysis. As additional extracted polymorphs’ structures, in situ ligand-bound complexes, seeded fibrils, and disease-associated fibrillar polymorphs whose structures will be solved in situ, become available, the pocket similarity landscape will need to be updated. This is not a weakness of the framework, but one of its main advantages: newly solved structures can be directly placed within the existing pocketome to determine whether they introduce novel binding environments or recapitulate already populated regions of amyloid pocket space.

Despite these limitations, the present study establishes a general framework for evaluating amyloid ligand targets in a comparative structural context. Rather than treating each fibril structure as an isolated object, the pocketome places every candidate binding site in relation to the full repertoire of known amyloid surface environments. This enables a more realistic assessment of which pockets are likely to support broad binding or genuine polymorph-restricted recognition.

### Concluding perspective

Overall, this work argues for a shift from a fold-centric to pocket-centric paradigm in amyloid ligand discovery. Fibril polymorphism remains essential for understanding amyloid biology and disease heterogeneity, but ligand selectivity is ultimately governed by the local binding environments exposed on fibril surfaces. By systematically mapping these environments across amyloid-β, tau, and α-synuclein fibrils, we show that many pockets are shared across proteins and polymorphs, whereas truly isolated pockets are rare.

This constrained pocket landscape helps explain why selective amyloid ligands have been difficult to obtain despite the growing availability of high-resolution fibril structures. It also provides a practical framework for future design efforts: prioritize pockets that are isolated in pocketome space, use *in vitro* fibrils when they reproduce the relevant local binding environment, and avoid recurrent cross-amyloid pockets that are structurally predisposed to off-target recognition. In this sense, the amyloid pocketome does not merely catalogue fibril cavities; it defines the structural limits and opportunities for selective amyloid targeting.

### Materials and Methods Creation of the dataset

Atomic structures of amyloid fibrils were retrieved from the Amyloid Atlas (81, 82) using Calypso scraping algorithm (52, 75). Only structures determined by cryo-electron microscopy (cryo-EM) were considered. ssNMR assemblies, and monomeric and peptide structures were excluded. Both apo fibrils and fibrils resolved in complex with small molecules were retained.

### Structural classification of fibril polymorphs

To classify fibril polymorphs independently of pocket analysis, pairwise structural comparisons were performed separately for α-synuclein, tau, and amyloid-β fibrils using the Calypso workflow from Connor et al. (52, 75). The resulting hierarchical classification was used to obtain polymorph groups by manually selecting a proper RMSD distance. The chosen distances were of 5.3, 4.8 and 3.3 Å for α-synuclein, tau and amyloid-β structures respectively. Polymorph group assignments were then used for comparative analysis with pocketome organization.

### Preparation of amyloid fibril structures

Each retained structure (polymorph group representatives and ligand-fibril complexes) was prepared using the Molecular Operating Environment (MOE) software (Chemical Computing Group). Preparation was performed using the QuickPrep protocol with default parameters. To ensure uniformity across the dataset, a maximum of four monomer layers was retained for each fibril, and no energy minimization or ligand refinement was applied to ligand-bound structures. Prepared structures were exported in MOL2 format for pocket detection.

### Detection and filtering of surface pockets

Surface pockets were detected on each prepared fibril structure using the VolSite algorithm implemented in the IChem toolkit. Pocket detection parameters were set according to those previously validated for protein–protein interaction cavities.

Detected pockets were filtered according to the following criteria:

1. Pocket volume must exceed 80 Å³.
2. Pockets located at the fibril tips (above the first or below the last retained monomer layer) were excluded.
3. Pockets located at the interface between protofilaments were excluded, as these regions lack sufficient accessibility for ligand binding.
4. Cavities buried within the fibril core or confined to narrow internal channels were excluded.
5. Hollow regions lacking well-defined residue contributions, as inferred from electron density maps, were excluded.

In fibrils composed of two identical protofilaments, symmetric copies of the same pocket were frequently detected. To avoid redundancy and prevent symmetry-driven bias in similarity calculations, only the larger of the two symmetric pockets was retained as a representative for subsequent analyses.

### Pocket descriptor calculation and similarity assessment

Each retained pocket was encoded using a set of 109 geometric and physicochemical descriptors, including volume, shape descriptors (e.g., asphericity, principal moments of inertia (73)), hydrophobicity, aromaticity, and electrostatic properties. Descriptor calculation followed previously established protocols. Pairwise pocket similarity was computed by calculating Euclidean distances between descriptor vectors using non-zero descriptor values only. Distances were normalized using a Gaussian kernel to yield the Pocket Similarity Index (PSI), defined as:

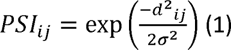

where *d_ij_* is the Euclidean distance between pockets *i* and *j*, and *σ* is the standard deviation of distances across all pocket pairs. PSI values range from 0 (maximally dissimilar) to 1 (identical). PSI matrices for global and protein-specific pocketomes are provided in the associated Zenodo repository.

### Pocketome visualization using minimum spanning trees

Pocketomes were visualized using a minimum spanning tree approach implemented through the TMAP framework. Starting from the complete graph defined by PSI matrices, edges were pruned to retain a spanning tree that maximizes total similarity while preserving connectivity among all pockets.

This representation enables global visualization of the pocket similarity landscape while maintaining local neighborhood relationships. Interactive HTML visualizations were generated for all pocketomes.

### Density-based clustering of pocket similarity space

To identify groups of highly similar and structurally isolated pockets, density-based spatial clustering (DBSCAN) was applied to PSI-derived distance matrices. Clustering was performed separately on global and protein-specific pocketomes.

DBSCAN parameters were selected to identify compact clusters characterized by high intra-cluster similarity and clear separation from surrounding pockets. Pockets not assigned to any cluster were considered part of the continuous similarity background. Cluster assignments were used exclusively for structural interpretation and do not imply functional annotation.

### Pocket annotation and visualization

Pocket residues were defined as amino-acid side chains contributing to cavity interactions as identified by VolSite. Residue-based annotations were used to label pockets according to sequence context and to identify recurrent residue-defined pocket types.

Pocket properties, cluster assignments, polymorph labels, and ligand-binding annotations were incorporated into interactive visualizations. All fibril structures, detected pockets, PSI matrices, and visualization files are freely available through the associated Zenodo repository (83).

## Acknowledgments

The authors thank Focus Biomarqueurs initiative from CEA and France Parkinson for funding G.O.; the Target ALS Foundation Inc. for funding Z.P.S.; and the Pasteur-Roux-Cantarini fellowship of Institut Pasteur for funding C.B.C.

