## Supplementary material for "A pocket-centric framework for selective targeting of amyloid fibril polymorphs": SI

**This PDF file includes:**

Figures S1 to S15

**Other supporting materials for this manuscript include the following:**

Supplementary Files 1 to 3

### Figures

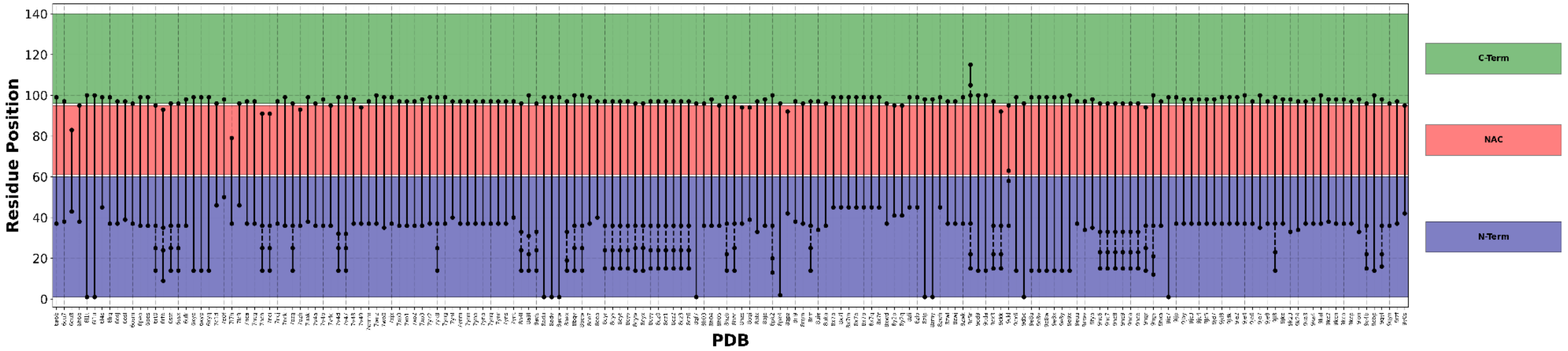


**Fig. S1:** Ordered residues from each α-synuclein structure retrieved from the Amyloid Atlas using Calypso. The residues are color-coded by alpha-synuclein domains. NAC = NonAmyloid Component.

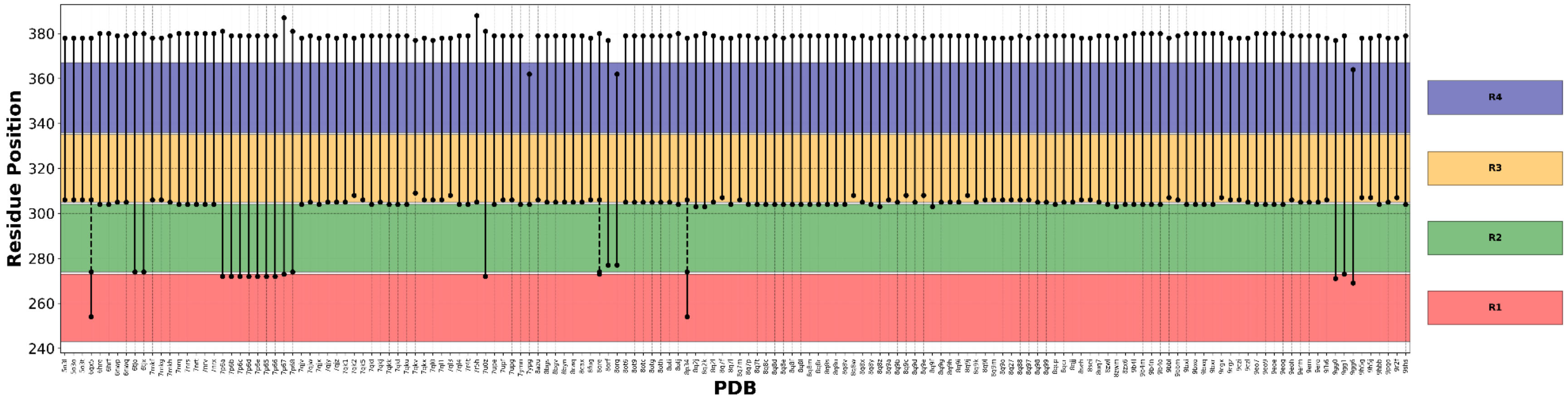


**Fig. S2:** Ordered residues from each tau structure retrieved from the Amyloid Atlas using Calypso. The residues are color-coded by tau repeat domains 1 to 4.

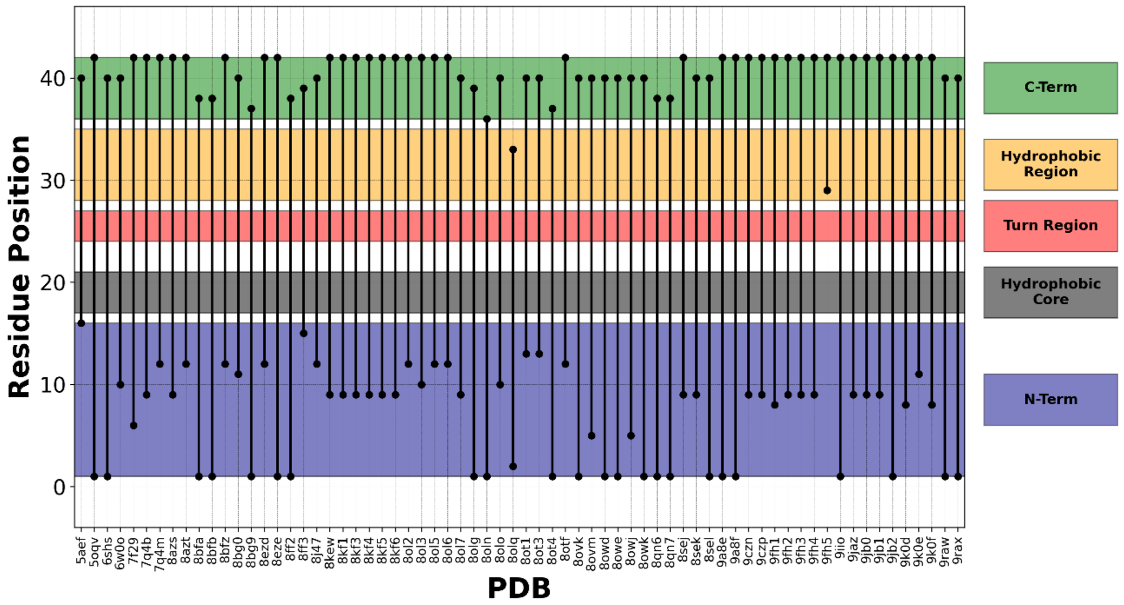


**Fig. S3:** Ordered residues from each amyloid-β structure retrieved from the Amyloid Atlas using Calypso. The residues are color-coded by protein domains.

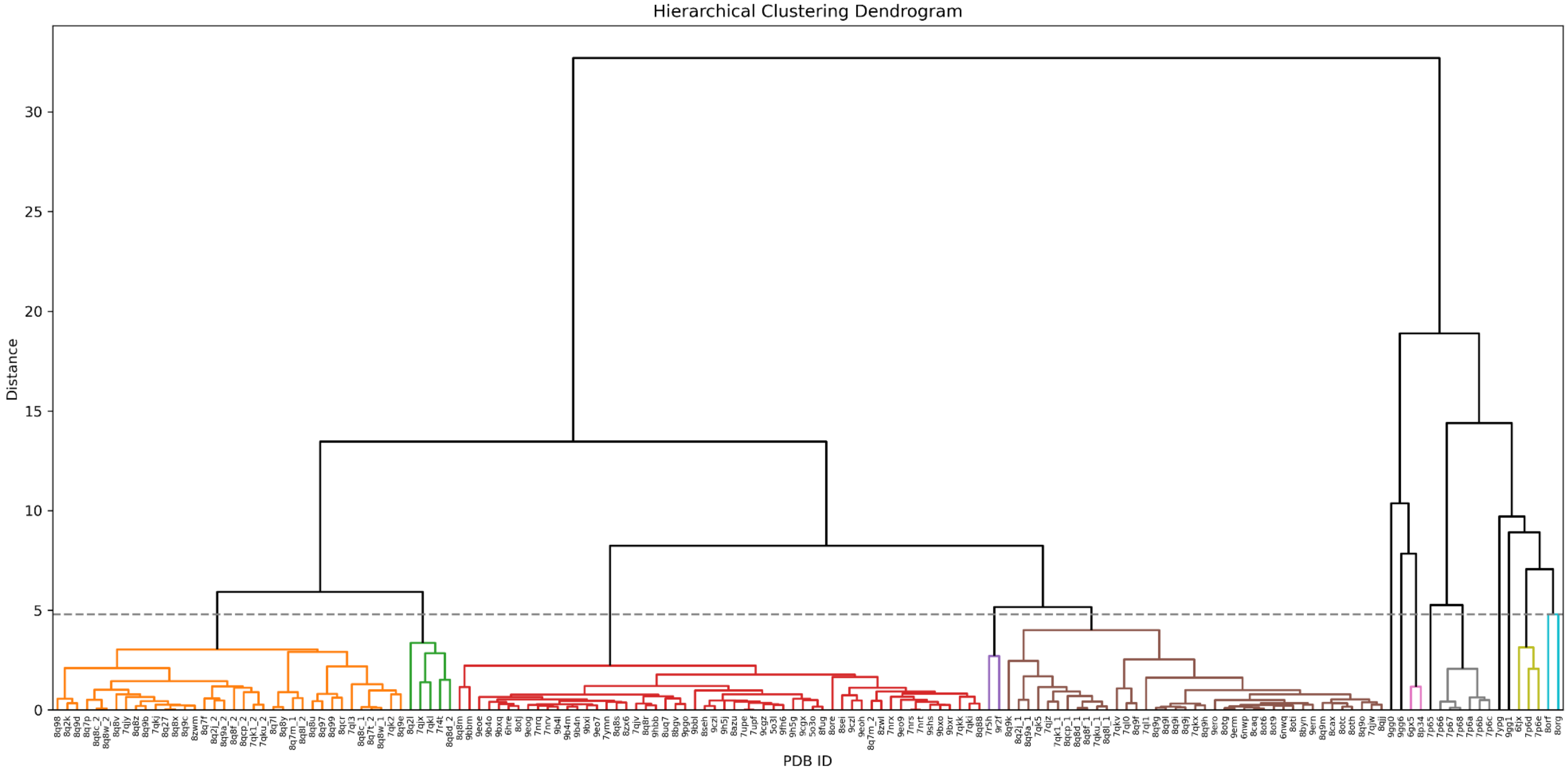


**Fig. S4:** Hierarchical classification of tau fibrils. The procedure was performed according to Connor et al. (1, 2). Fibrils were classified into 14 groups, cutting at a distance of 4.8 Å. Fibrils from same groups are colored accordingly. Although determined as a singlet structure, PDB ID 9GG6 was not conserved for further analysis and assimilated with results from PDB 9GG0 and 6GX5.

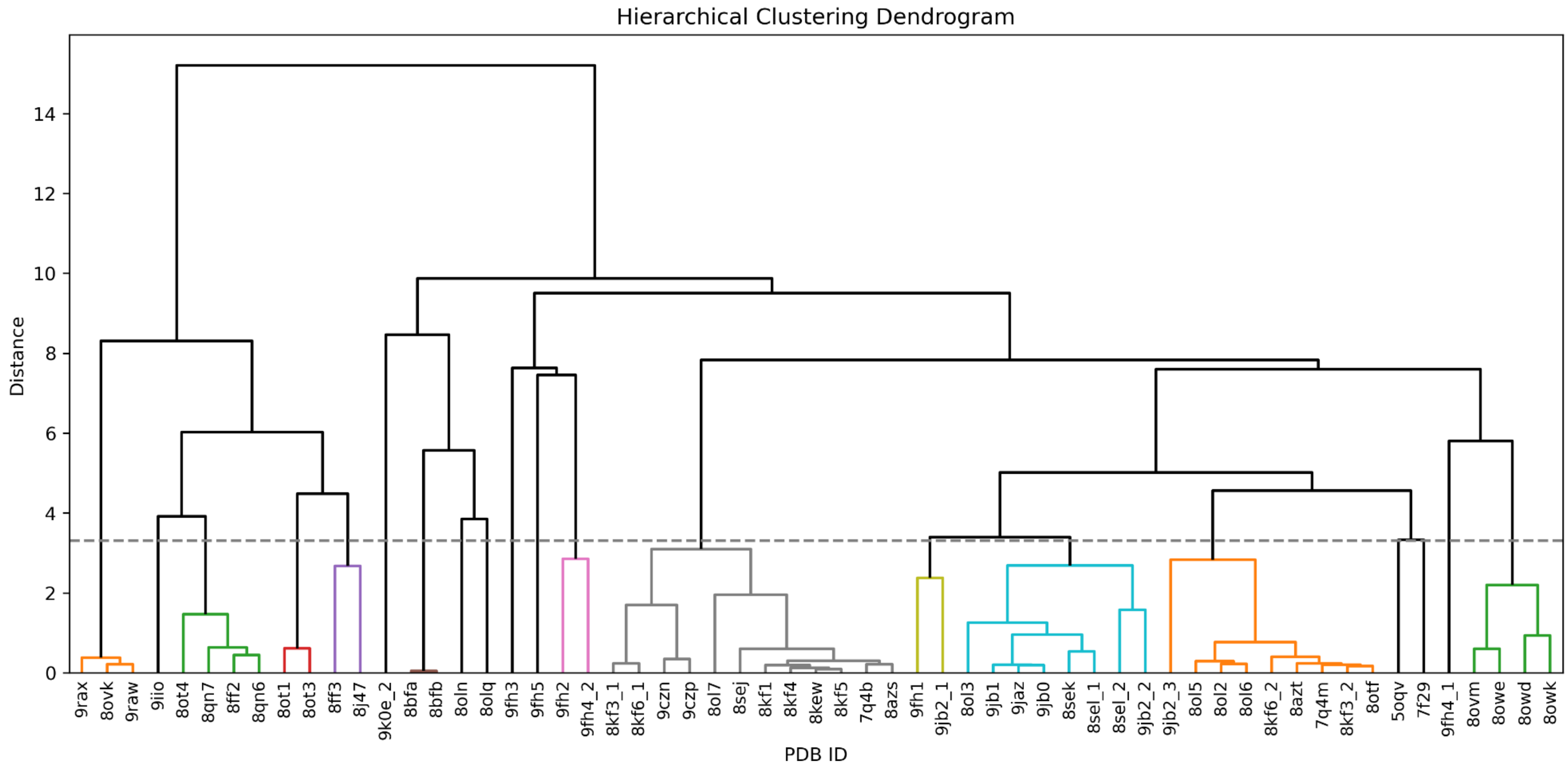


**Fig. S5.** Hierarchical classification of amyloid-β fibrils. The procedure was performed according to Connor et al. Fibrils were classified into 20 groups, cutting at a distance of 3.3 Å. Fibrils from same groups are colored accordingly.

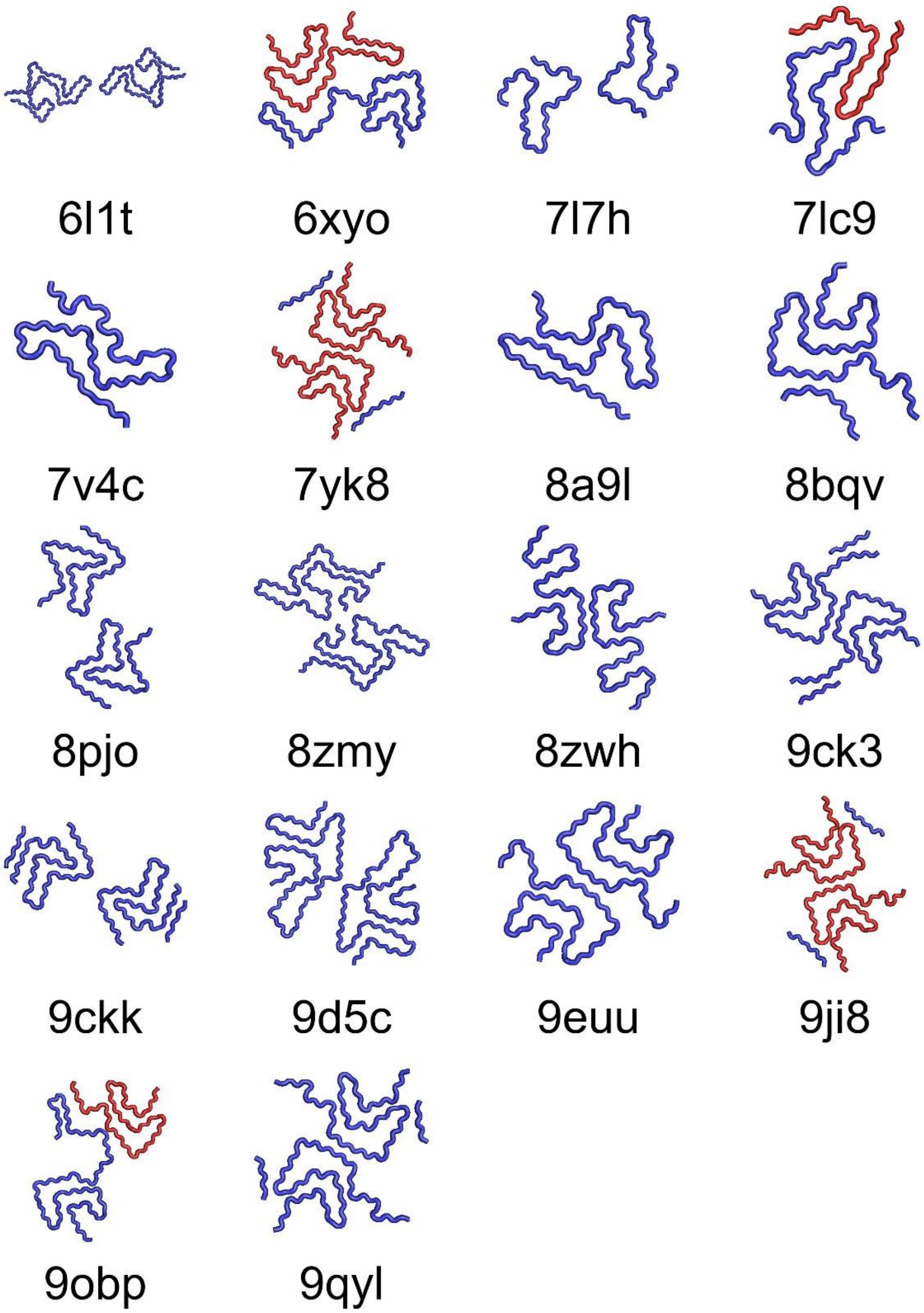


**Fig. S6:** Representative folds for each α-synuclein polymorph group identified after hierarchical classification. Structures are color-coded by asymmetric unit, with unit number 1 in blue and number 2 in red. The representative unit considered in the study were always number 1. PDB IDs 7YK8 and 9JI8 were removed from the study, as identified as singlets through the additional unresolved peptide.

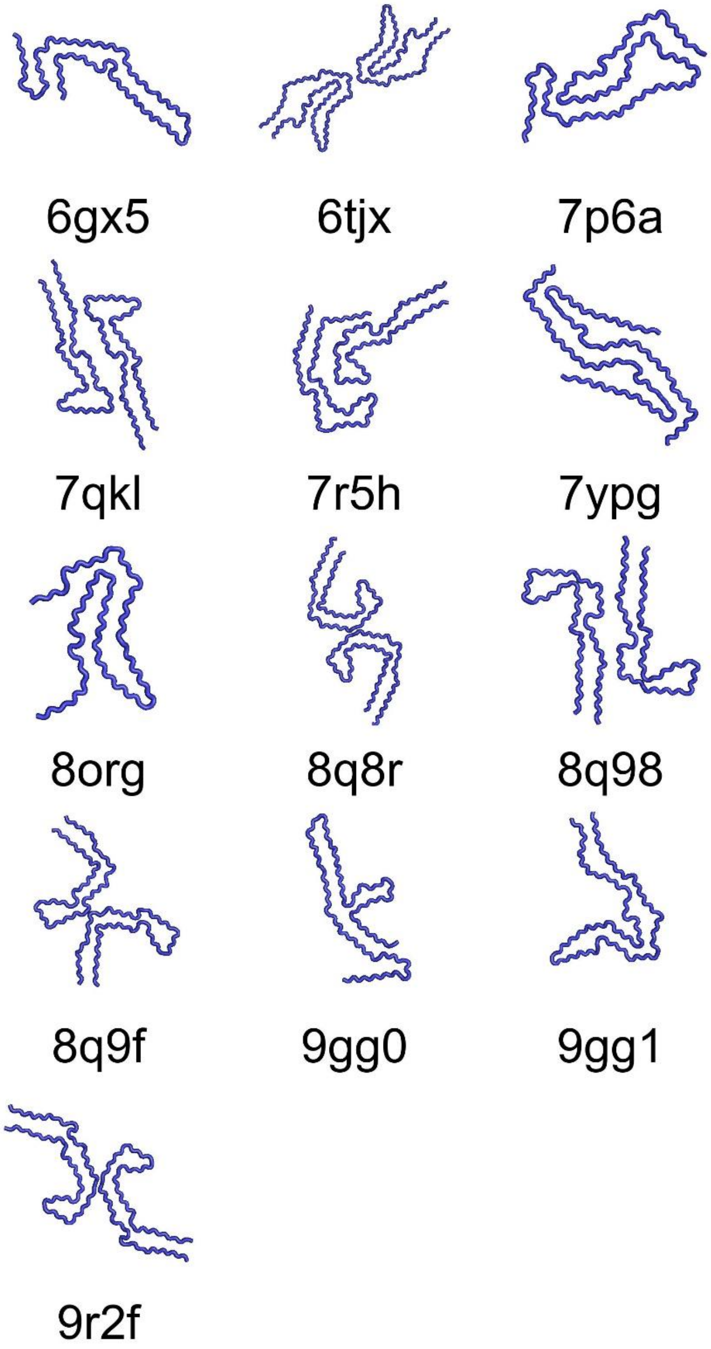


**Fig. S7:** Representative folds for each tau polymorph group identified after hierarchical classification.

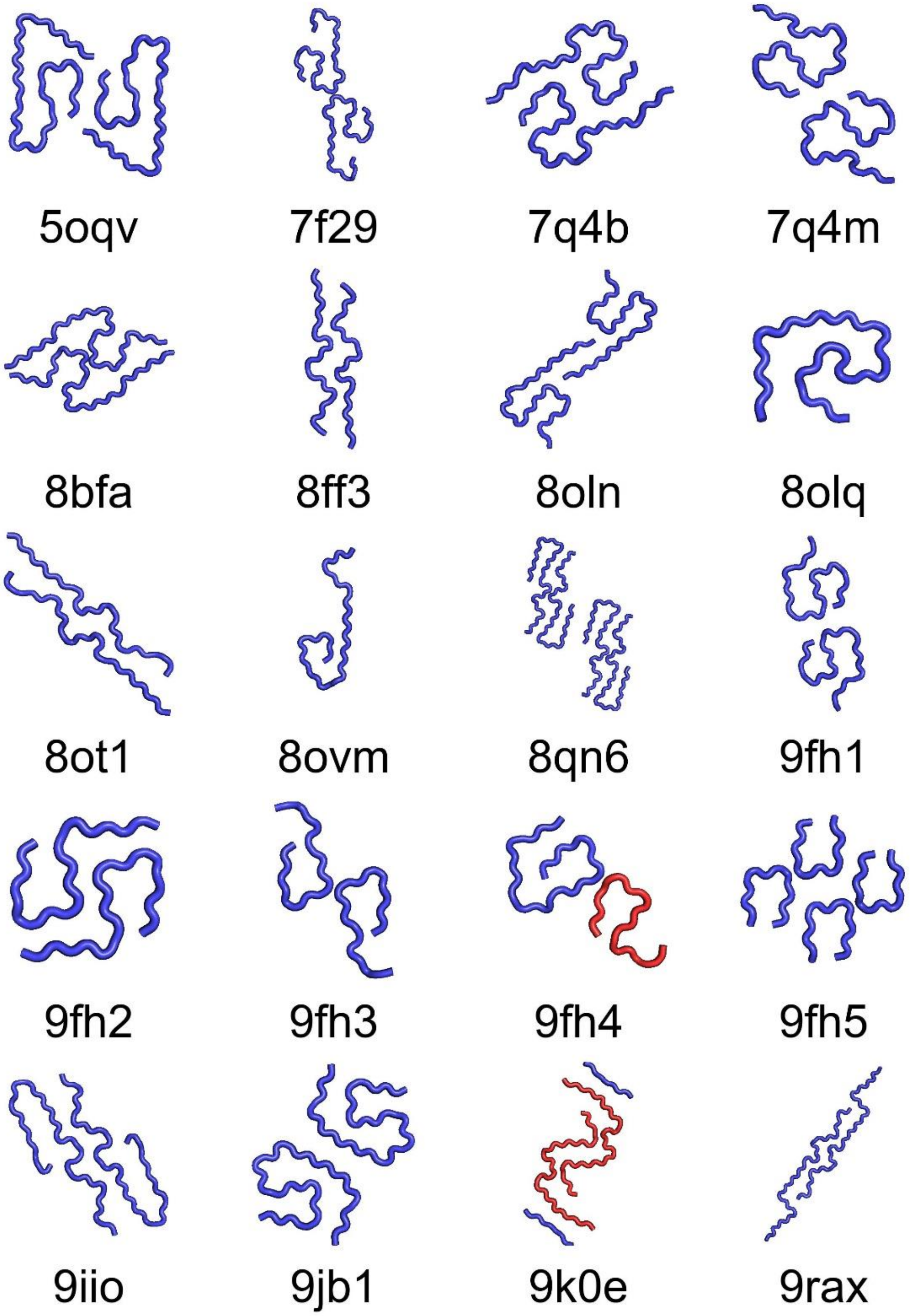


**Fig. S8:** Representative folds for each amyloid-β polymorph group identified after hierarchical classification. Structures are color-coded by asymmetric unit, with unit number 1 in blue and number 2 in red. The representative unit considered in the study were always number 1.

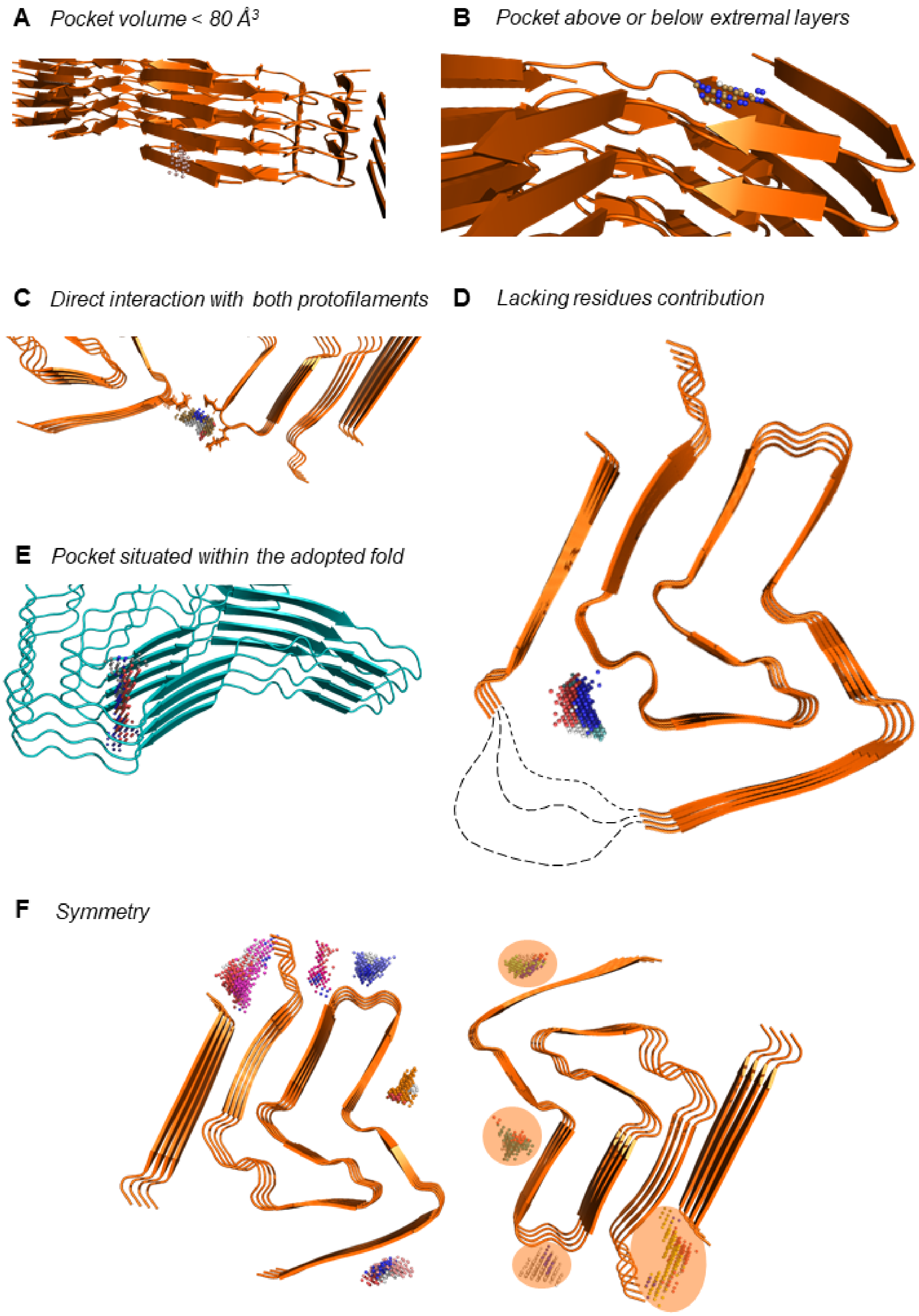
A Pocket volume < 80 Å3
B Pocket above or below extremal layers
C Direct interaction with both protofilaments
D Lacking residues contribution
E Pocket situated within the adopted fold
F Symmetry

**Fig. S9:** Overview of the different exclusion criteria considered in our study. Fibril backbones are represented in orange (for α-synuclein (αSyn) fibril polymorph 2a (PDB: 6SSX), used across all shown examples) or blue (for tau monofilament extracted from corticobasal degeneration (CBD) brain tissue (6TJO)) depending on the protein, and pockets as dense point clouds with several colors. A) Pocket volumes must be greater than 80 $Å^3$. Example given is Cavity_N45 which volume is 30 $Å^3$. We can see that the pocket has only been found by VolSite on two monomer layers out of the four considered. B) Pockets must not be located at the fibril extremities. The given example shows Cavity_N42, found outside of the four extremal layers. C) The pocket must not interact directly with both protofilaments. Example of cavity_N3, with highlighted side chain of interacting residues from both protofilaments (E57, K58 and K45'). Such interaction is prohibited since the energy of the ligand binding to the pocket could destabilize the whole fibril or not overcome the saline intramolecular interaction and thus be unable to bind favorably. D) The pockets must not be localized in a space lacking the contribution of key residues. Example given is cavity_N4, found between residues G25 and G36. Hypothetic possible lacking sequence structures are drawn in dashed lines. E) The pocket must not be localized within the adopted monomeric fold. The shown example is of cavities_N10, 12 and 14 from CBD monofilament that together form a cavity found in a narrow space inside the adopted fold, where even water molecules are unable to access. F) Pockets considered from the αSyn fibril 6SSX after meeting all previously mentioned exclusion criteria. The redundant pockets found across both protofilaments are highlighted with orange transparent oval shapes. The symmetry filter retains the biggest pocket from the two.

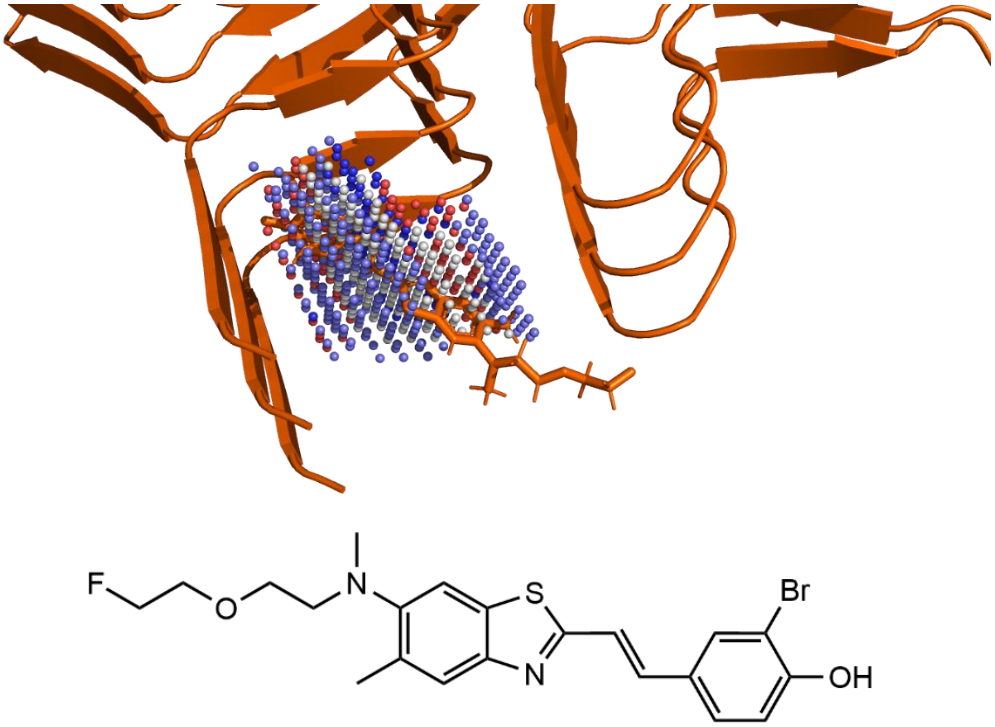


**Fig. S10:** Detection of pockets in the ligand space from ligand-amyloid fibril complexes files. Top: Close-up view of the Cavity_N1 (511 Å$^3$) from αSyn *in vitro* fibil-F0502B ligand complex (PDB: 7WMM). We can see that the cavity matches the core benzothiazole-ethenyl-phenol structure of the ligand, as described by the authors after cryo-EM reconstruction. Bottom: Molecular structure of binding ligand F0502B.

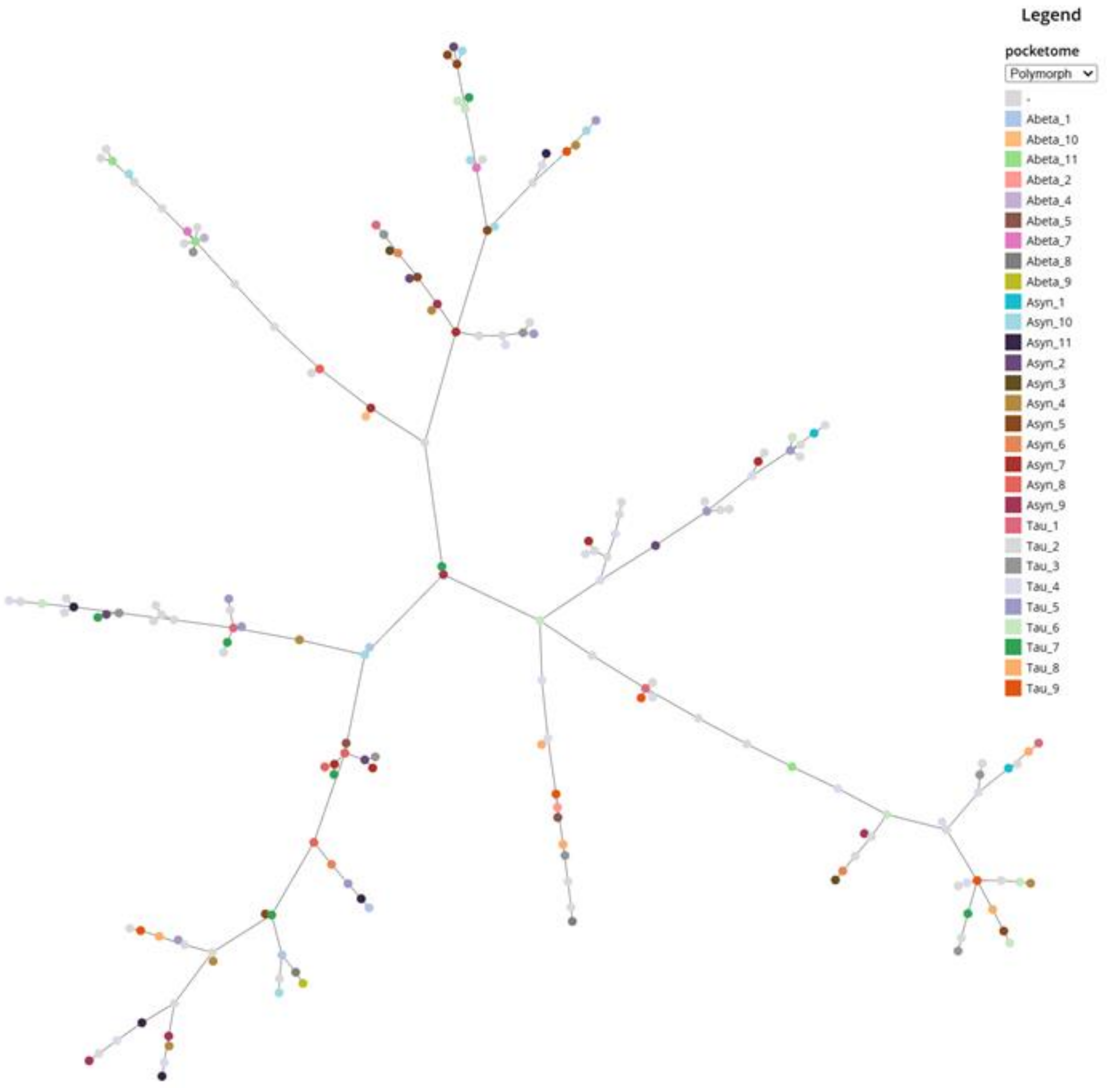


**Fig. S11:** The amyloid pocketome, represented as a Tree map, and colored by identified polymorph groups after hierarchical classification. Each pocket is represented using a dot, and dots are connected using the PSI value, so that the total number of branches is minimal. The legend shown here represents identified polymorph groups.

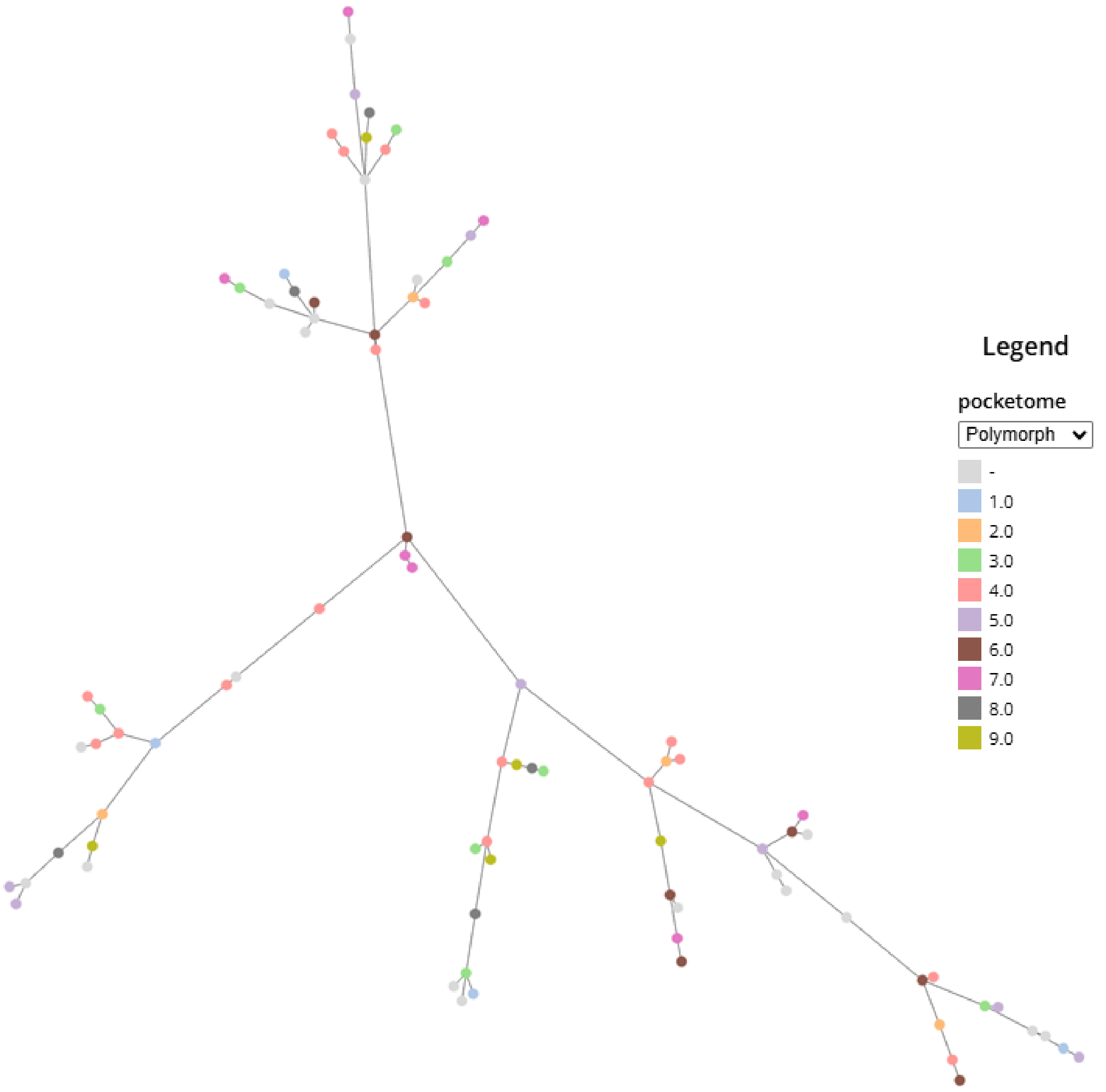


**Fig. S12:** The tau pocketome, colored by polymorph group. Each pocket is represented using a dot, and dots are connected using the PSI value, so that the total number of branches is minimal. The legend shown here represents identified polymorph groups.

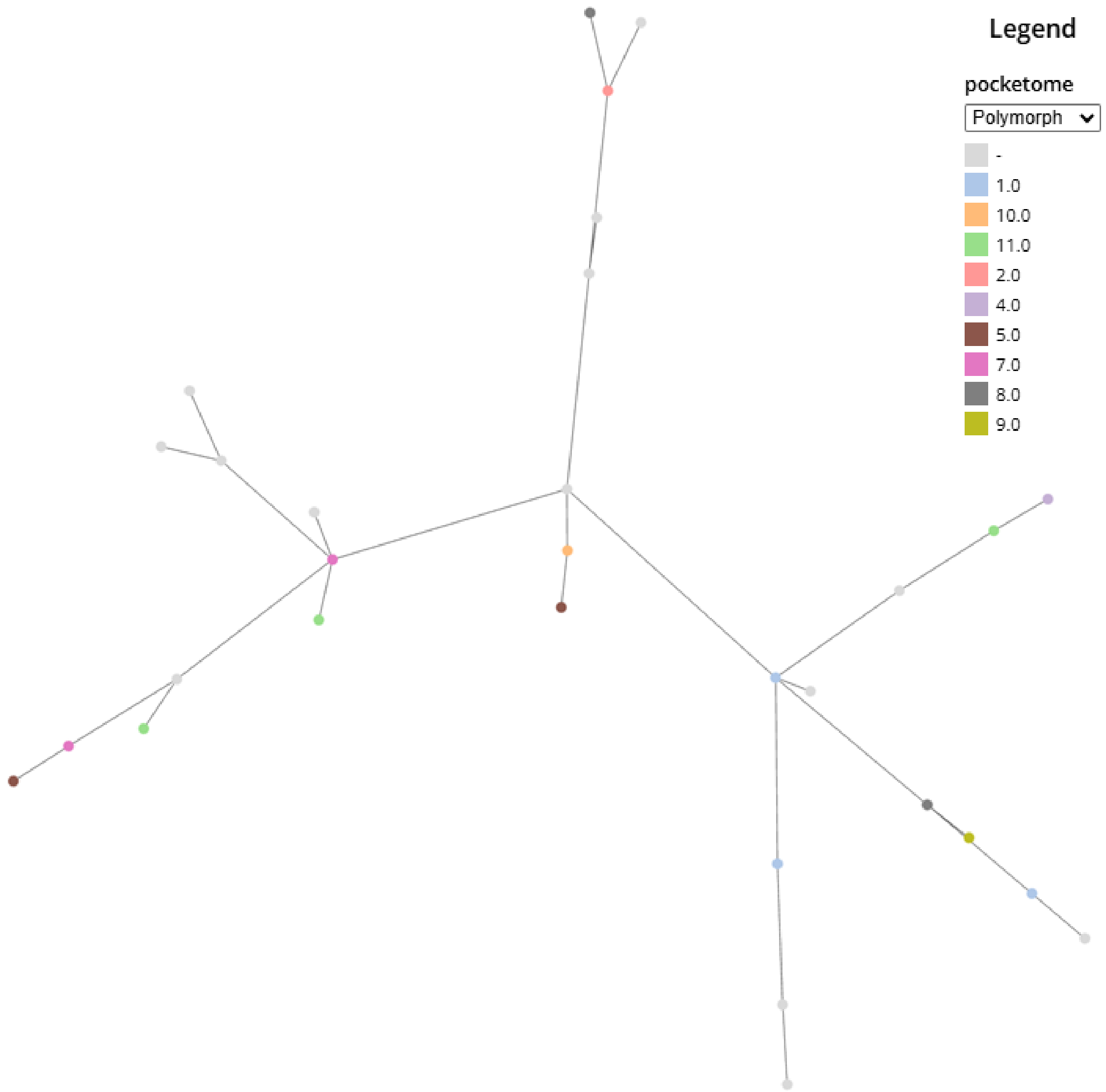


**Fig. S13:** The amyloid-beta pocketome, colored by polymorph group. Each pocket is represented using a dot, and dots are connected using the PSI value, so that the total number of branches is minimal. The legend shown here represents identified polymorph groups.

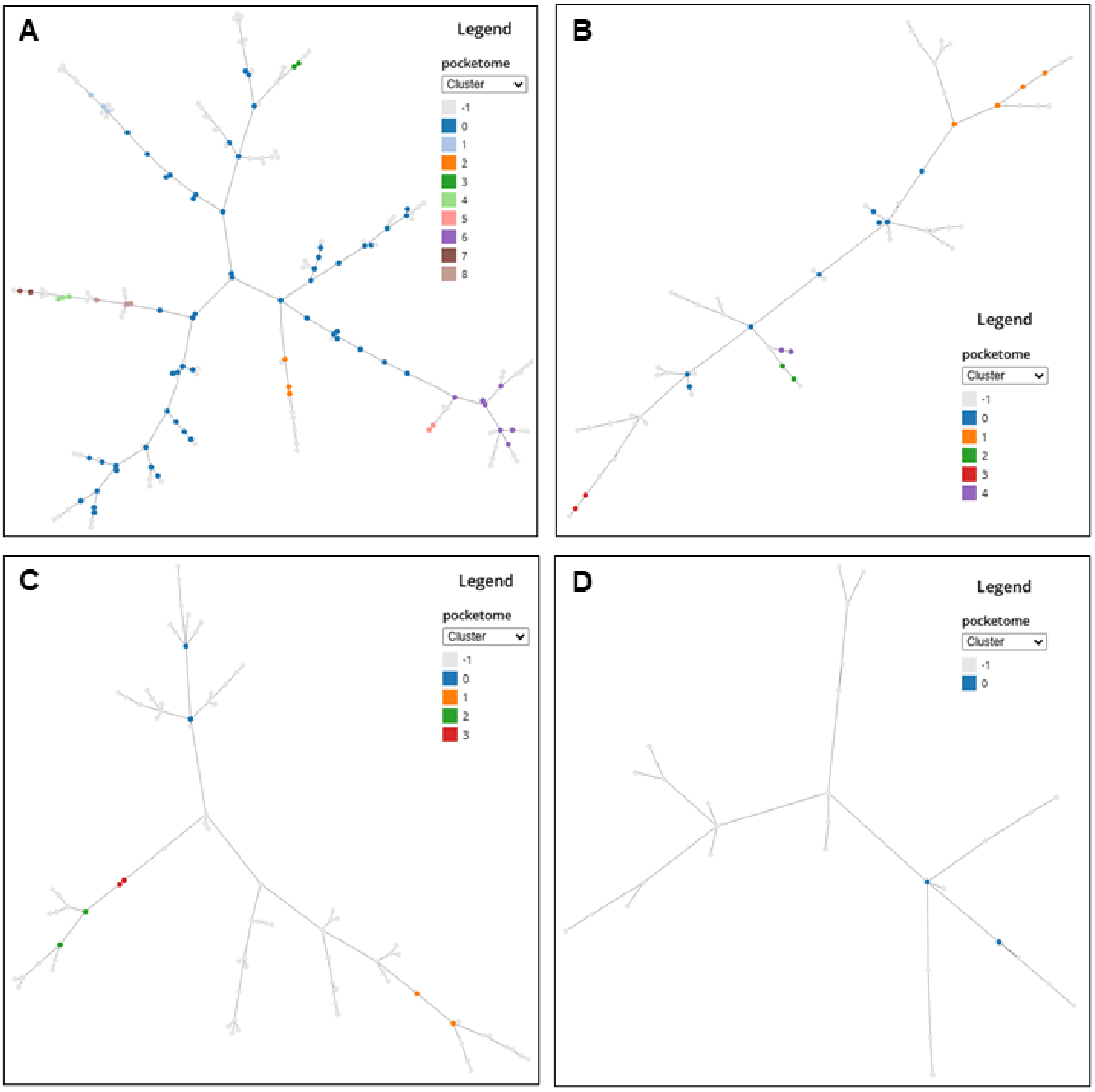


**Fig. S14:** The amyloid and individual pocketomes, colored after density-based clustering of each dataset. Each pocket is represented using a dot, and dots are connected using the PSI value, so that the total number of branches is minimal. The legend shown here represents determined clusters. Clustered pockets are colored accordingly, and non-clustered pockets are colored in grey and labeled as "-1". A) The amyloid pocketome. B) Tha alpha-synuclein pocketome. C) The tau pocketome. D) The amyloid-beta pocketome.

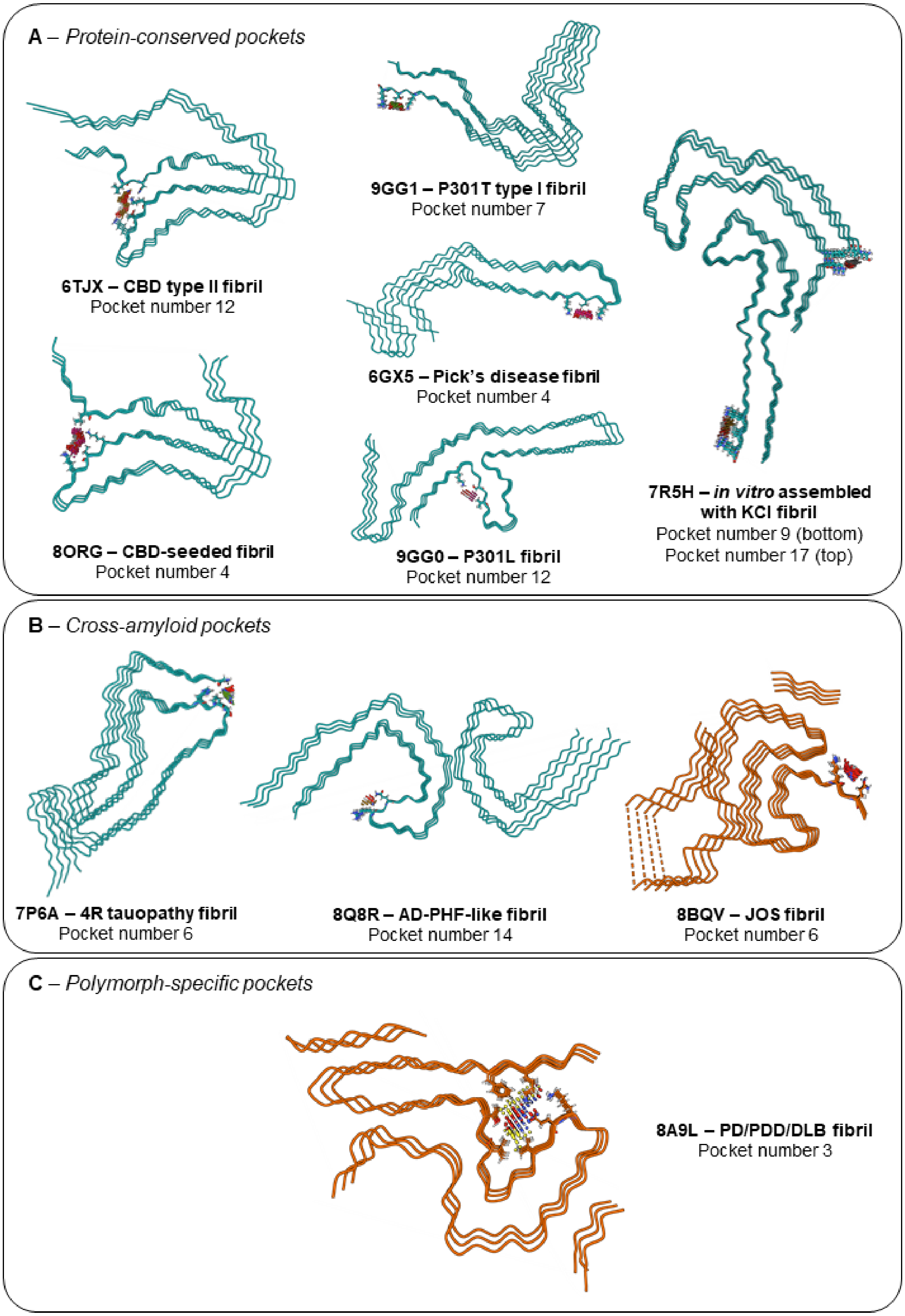
A – *Protein-conserved pockets*
6TJX – CBD type II fibril
Pocket number 12
8ORG – CBD-seeded fibril
Pocket number 4
9GG1 – P301T type I fibril
Pocket number 7
6GX5 – Pick's disease fibril
Pocket number 4
9GG0 – P301L fibril
Pocket number 12
7R5H – *in vitro* assembled with KCl fibril
Pocket number 9 (bottom)
Pocket number 17 (top)
B – *Cross-amyloid pockets*
7P6A – 4R tauopathy fibril
Pocket number 6
8Q8R – AD-PHF-like fibril
Pocket number 14
8BQV – JOS fibril
Pocket number 6
C – *Polymorph-specific pockets*
8A9L – PD/PDD/DLB fibril
Pocket number 3

**Fig. S15:** Illustrative examples from cluster analysis. A) Pockets from amyloid cluster number 6 from tau structures, constituting an example of protein-conserved pockets. Fibril backbones are represented in blue and pockets as dense point clouds. B) Pockets from amyloid cluster number 4 from alpha-synuclein and tau structures, constituting an example of cross-amyloid pockets. Alpha-synuclein is shown in orange and tau in blue, with pockets as dense point clouds. C) Alpha synuclein extracted fibril from Parkinson's Derived Dementia brain tissue and detected pocket number 3. The surrounding and interacting residues' side chains are highlighted. This pocket was found to have a PSI value of 0.005 with its first neighbor.
